# BIRC3: A Prognostic Predictor and Novel Therapeutic Target in TMZ-Resistant Glioblastoma Tumors

**DOI:** 10.1101/2023.08.23.554432

**Authors:** M Morelli, S Franceschi, F Lessi, P Aretini, A Pastore, E Corradi, A Marranci, C. Gambacciani, F Pieri, G. Grimod, N Montemurro, M Giacomarra, M Menicagli, G Ferri, Francesco Pasqualetti, M Sanson, A Picca, AL Di Stefano, OS Santonocito, CM Mazzanti

## Abstract

**Background:** Glioblastoma (GB) is an incurable malignant tumor of the central nervous system, with a poor prognosis. Robust molecular biomarkers associated with therapeutic response or survival are still lacking in GB. Previously, using NADH-fluorescence lifetime imaging (NADH-FLIM), as a new drug screening precision medicine ex-vivo approach, we categorized patient-derived vital tumors into TMZ responder (Resp) and non-responder (Non-Resp) groups, revealing differentially expressed genes.

**Methods:** Expanding on our previous study, we assessed TMZ response in a larger cohort of primary and recurrent ex-vivo live GB tumors (n=33) using NADH-FLIM. Transcriptome analysis was performed to characterize TMZ Resp and Non-Resp cases, and in-silico and functional cellular investigations were conducted to explore the efficacy of potential biomarkers.

**Results:** Genes dysregulated in the previous study showed consistent expression patterns. BIRC3, a potent apoptosis inhibitor, was significantly upregulated in TMZ-resistant samples. BIRC3 expression complemented MGMT status as a prognostic factor in multiple TCGA cohorts. BIRC3 functioned as a prognostic factor of survival also in separate European private glioblastoma cohorts. The BIRC3 antagonist, AZD5582, in combination with TMZ, effectively reversed TMZ resistance by restoring apoptosis in glioblastoma cell lines and patient-derived organoids.

**Conclusions:** BIRC3 holds promise as a prognostic biomarker and predictor of TMZ response in GB. Assessing BIRC3 expression could aid in stratifying patients for combined TMZ and AZD5582 therapy. Our study highlights the potential of functional precision medicine and BIRC3 assessment as a standard tool in glioblastoma clinical oncology, improving outcomes.

**KEYPOINTS:**

- BIRC3, previously overlooked, identified through dynamic precision medicine using TMZ perturbation of glioblastoma tissue as a robust prognostic factor.
- The gene BIRC3 is an independent prognostic factor associated with shorter survival and TMZ resistance, rigorously validated across various case studies and datasets, including two expansive European case studies.
- Proposal of anti-BIRC3 drug, AZD5582, shows promise as a novel therapeutic option to overcome TMZ resistance in GB tumors, providing hope for improved outcomes and personalized treatment strategies for patients with limited treatment options

**IMPORTANCE OF THE STUDY:** Glioblastoma (GB), an aggressive cancer type with a bleak prognosis, lacks dependable biomarkers for treatment prediction. Few markers like MGMT promoter methylation, IDH1 mutation, TERT gene mutations, and EGFR amplification are known, but their predictive consistency varies. Temozolomide (TMZ) resistance, seen in over 50% of GB patients, complicates matters. BIRC3, an apoptosis-inhibiting gene, displays heightened expression in TMZ-resistant tumors. Our study examined BIRC3 in GB patient samples, finding it an independent prognostic factor linked to shorter survival and TMZ resistance. Our research builds upon Wang et al.’s 2016 and 2017 findings, delving deeper through TCGA data and European case studies. BIRC3’s consistent prominence suggests its significance, with functional experiments confirming its role. We assessed AZD5582, targeting BIRC3, which, when combined with TMZ, curtailed cell growth and induced apoptosis. Notably, AZD5582 countered TMZ resistance in patient-derived GB-EPXs, except for low BIRC3 cases. Our precision medicine approach enhances personalized therapies and outcomes, highlighting BIRC3’s potential as a prognostic marker and AZD5582 as a new therapy for TMZ-resistant GB.

## INTRODUCTION

Glioblastomas (GBM) are incurable brain tumors; despite aggressive treatment, consisting of surgical resection, chemotherapy, and radiotherapy, the median survival is 15 months [1]. After decades of intense research into the biology and treatment of GBM, overall survival remains stagnant, with very few approved chemotherapeutic or biological agents. It is now accepted that glioblastoma is a heterogeneous fatal disease and that different patients’ tumors present different genetic, transcriptomic, and epigenetic footprints with important implications on their evolution and progression, and therefore on clinical recommendations [2]. The 2016 WHO classification of the central nervous system (CNS) tumors broke the principle of diagnosis based entirely on histological features and introduced genetic parameters into the classification of CNS tumors. Exploration of novel and reliable biomarkers for the prediction of glioblastoma prognosis may help to elucidate the molecular mechanism of glioblastoma development and progression [3]. Molecular profiling has provided clinical benefits for patients with various advanced cancer [4, 5]. Understanding the mutational profile of a tumor is important to inform diagnosis and guide therapeutic options, as well as identify patients that may not respond to treatment. Multiple three-dimensional (3D) tumour organoid models assisted by multi-omics and Artificial Intelligence (AI) have contributed greatly to preclinical drug development and precision medicine[6]. The key advantage of precision medicine in oncology is the ability to match the right drug to the right patient to help improve patient outcomes. In the management of glioblastoma patients, in addition to clinical factors such as age, performance status, and extent of resection, only one specific molecular alteration is prognostic and to some degree predictive of treatment response: the level of methylation of the promoter region of the DNA repair enzyme methylguanine methyltransferase (MGMT) [7]. Approximately 30% of tumors have MGMT promoter methylation, and those patients respond better to treatment and live longer than patients with unmethylated MGMT [5, 8]. In the absence of promoter methylation, MGMT enzyme transcription is uninterrupted thereby resulting in the repair of TMZ-induced DNA alkylating events. Nonetheless, even patients with positive MGMT methylation status will invariably fail TMZ treatment, suggesting that there are potentially alternate mechanisms for TMZ drug resistance independent of methylation status [9]. Unfortunately, aside from MGMT promoter methylation and TMZ response, molecular biomarkers associated with therapeutic response or prognostic of survival are still lacking in GBM. Previously in a recent study, using NAD(P)H - fluorescence lifetime imaging (NAD(P)H -FLIM), as a new drug screening precision medicine ex-vivo approach, we perturbed GBM patients-derived vital tumor explants (GB-EXPs) by TMZ treatment and obtained a stratification in TMZ responder (Resp) and non-responder (Non-Resp) tumors, identifying distinct differentially expressed genes [10, 11]. Several of the identified genes resulted already associated with glioblastoma and response to TMZ reinforcing our novel functional precision medicine approach. Among the various differentially expressed genes, we focused our attention on the BIRC3 gene that we found significantly upregulated in TMZ-resistant tumors [10]. BIRC3 codes for the Baculoviral IAP repeat-containing protein (also known as cIAP2) that is a member of the inhibitor of apoptosis family (IAP) that suppresses apoptosis by interfering with the activation of caspases [12, 13]. Recently Wang et al [14] has involved BIRC3 in a novel mechanism of TMZ drug resistance resulting in apoptosis evasion in GBM. In particular, their work for the first time shows that TMZ and RT treatment induce BIRC3 gene expression up-regulation. This resulting up-regulation of BIRC3 confers onto human GBM cells as well as human GBM xenografts an apoptotic evasion phenotype [14].

In this study, we aimed to expand upon our previous glioblastoma case study by employing the NAD(P)H -FLIM metabolic approach to assess the response of primary and recurrent live GB-EXPs to TMZ treatment. Additionally, we conducted transcriptome analysis to molecularly characterize cases that were responsive or non-responsive to TMZ. Consistent with our earlier findings, we observed the same directional dysregulation of gene expression in the majority of genes and confirmed the statistically significant upregulation of BIRC3 in TMZ-resistant samples. assess the BIRC3 mRNA expression and its complementarity with the MGMT status as a predictor of outcome in several cohorts analyzed with different RNA analysis approaches in the large public TCGA data set. To confirm in silico results we proved that BIRC3 functioned as a prognostic factor of survival also in a French and Italian private glioblastoma cohort analyzed by us through RNAseq and RT-PCR, respectively. In addition, considering that a major mechanism of drug resistance lies in the dysfunctional survival signaling in cancer cells and the reluctance of these cells to undergo apoptosis, drug agents that can restore sensitivity of these cells were taken into consideration. Notably, we investigated the effects of AZD5582, an antagonist of BIRC3 developed in 2015[15], either as a standalone treatment or in combination with TMZ. Our results indicate that AZD5582 treatment has the potential to overcome TMZ resistance and suggests that BIRC3 could serve as a promising therapeutic target in glioblastoma tumors that are unresponsive to standard TMZ therapy.

## 1) MATERIALS AND METHODS

### SAMPLES

#### Italian samples

The study was performed in accordance with the Declaration of Helsinki and the sample collection protocol was approved by the Ethics Committee of the University Hospital of Pisa (787/2015). Tumors were obtained from patients who had undergone surgical resection of histologically confirmed GBM after obtaining informed consent. Samples were obtained from the Neurosurgery Department the Unit of Neurosurgery of Livorno Civil Hospital Italian (Table 1, Table S7). The patients clinical and demographic data are presented in Table 1. All cases had a diagnosis of GB with no previous history of any brain neoplasia and were not carrying R132 IDH1 or R172 IDH2 mutations. Details of samples handling procedures are available in supplementary materials. *French samples*: The patient cohort includes retrospective6 and prospective cases of gliomas identified (since January 2014) at Pitié-Salpêtrière Hospital. Tumor specimens, blood samples, and clinico-pathological information were collected with informed consent and relevant ethical board approval in accordance with the tenets of the Declaration of Helsinki. For the samples, clinical data and follow-up are available in the neuro-oncology database (OncoNeuroTek, GH Pitié-Salpêtrière, Paris).

**Table 1:** GB-EXPs NADH-FLIM stratification of TMZ Responder (Resp) and Non Responder (Non-Resp)

| GB code | Sex | Age | Primary/<br>Recurrence | NADH<br>FLIM<br>TMZ<br>Response | % of Drug<br>Response | Idh1/2<br>molecular<br>status | MGMT<br>methylat<br>ion |
| --- | --- | --- | --- | --- | --- | --- | --- |
| GB 1 | m | 59 | P | Resp | MR | WT | unmeth |
| GB 2 | m | 57 | P | Resp | MR | WT | meth |
| GB 3 | m | 30 | P | Resp | MR | WT | unmeth |
| GB 4 | f | 46 | P | Resp | MR | WT | meth |
| GB 5 | m | 71 | P | Resp | LR | WT | meth |
| GB 6 | m | 65 | P | Resp | HR | WT | meth |
| GB 7 | m | 80 | P | Resp | HR | WT | meth |
| GB 8 | f | 59 | P | Non-Resp | NR | WT | unmeth |
| GB 9 | f | 59 | P | Non-Resp | NR | WT | unmeth |
| GB 10 | f | 59 | P | Non-Resp | NR | WT | unmeth |
| GB 11 | m | 72 | P | Non-Resp | NR | WT | unmeth |
| GB 12 | f | 40 | P | Non-Resp | NR | WT | meth |
| GB 13 | f | 73 | P | Non-Resp | NR | WT | unmeth |
| GB 14 | m | 47 | P | Non-Resp | NR | WT | unmeth |
| GB 15 | f | 68 | P | Resp | LR | WT | meth |
| GB 16 | f | 59 | R | Resp | LR | WT | unmeth |
| GB 17 | f | 68 | P | Non-Resp | NR | WT | meth |
| GB 18 | m | 72 | P | Non-Resp | NR | WT | unmeth |
| GB 19 | m | 72 | P | Resp | HR | WT | meth |
| GB 20 | m | 55 | P | Resp | HR | WT | meth |
| GB 21 | f | 74 | P | Non-Resp | NR | WT | meth |
| GB 22 | m | 76 | P | Non-Resp | NR | WT | meth |
| GB 23 | m | 73 | R | Resp | HR | WT | meth |
| GB 24 | m | 73 | R | Resp | HR | WT |  |
| GB 25 | f | 75 | R | Resp | NR | WT |  |
| GB 26 | m | 73 | R | Resp | LR | WT | meth |
| GB 27 | m | 73 | R | Resp | LR | WT | meth |
| GB 28 | f | 69 | R | Non-Resp | NR | WT | meth |
| GB 29 | f | 57 | R | Non-Resp | NR | WT | unmeth |
| GB 30 | m | 56 | R | Non-Resp | NR | WT | unmeth |
| GB 31 | m | 70 | R | Non-Resp | NR | WT | unmeth |
| GB 32 | m | 70 | R | Non-Resp | NR | WT | meth |
| GB 33 | m | 57 | R | Non-Resp | NR | WT | unmeth |

#### GBM commercial cell lines

T98G GBM cell lines were obtained from American Type Culture Collection (ATCC, Rockville, MD). T98G were grown as monolayers in Dulbecco’s Modified Eagle Medium (DMEM) low glucose and high glucose respectively without red phenol, supplemented with 10% FBS and 1% Penicillin-Streptomycin. For FLIM experiments, cells were grown in a 35 mm Nunc Glass Bottom Dishes (Thermo Fisher Scientific).

### RNA EXPRESSION ANALYSES

#### RNA extraction

RNA was extracted from tumor samples, and cells using Maxwell 16 LEV Simply RNA Tissue Kit (Promega, Madison, WI, USA), according manufacturer’s protocol. RNA was tested using the Qubit Fluorometer (Life Technologies, Carlsbad, CA) and Agilent 2200 Tapestation (Agilent Technologies, Santa Clara, CA) system. NGS library preparation details are available in supplementary materials.

#### BIRC3 Real time PCR

cDNA was synthesized starting from 250 ng of total RNA using iScript™ cDNA Synthesis Kit (Bio-Rad) and real-time PCR was performed following the manufacturer’s protocol of Universal SYBR Green Supermix (Bio-Rad). Ct values were obtained for each gene and normalized to β-actin.. The following primers were used: Birc3 (F: 5′-GATCCATGGGTTCAA -3′; R: 5′-GGCTTGAACTTGACG -3′), β-actin (F: 5′-CTGGCACCACACCTTCTAGA-3′; R: 5′-GTACATGGCTGGGGTGTTGA -3′). Birc3 protein expression analyses by IHC, IC and western blot are available in supplementary materials.

#### FLOW CYTOMETRY-Apoptotic Assay

A total of 100000 cells per well were seeded in a 6-well plate format and treated with 500 uM TMZ, 5 uM AZD, and a combination of the two drugs. After 72 h, the stage of apoptosis was assessed using the Annexin V FITC kit (Beckman Coulter, IM2375) according to the manufacturer’s instructions. The percentages of cells in the different stages of apoptosis were detected by using the CytoFLEX S flow cytometer (Beckman Coulter).

#### GB-Explants culturing, NADH-FLIM data acquisition and data analysis

All details have been reported in previous papers and are available in supplementary materials [10, 16].

#### TMZ and AZD5582 TREATMENTS

The drug used in this study was TMZ and AZD5582 (Sigma, St. Louis, MI, USA). TMZ was dissolved in DMSO to prepare a stock concentration of 100 mM, and then, diluted to the required concentrations with complete cell culture medium. AZD5582 was dissolved in water to reach a stock concentration of 4,9 mM. T98G cell line was treated at 30% confluency, replacing medium with fresh one, containing TMZ and AZD5582 a different doses for treated cells, and an equal volume of DMSO and sterile water for controls. Cultures were exposed to both treatments for 72hr. GB-EXPs both matrigel 3 days after being cultured, were treated with TMZ and AZD5582. Medium volume was replaced with fresh medium containing TMZ 600 μM, 25uM and 50uM AZD5582, alone, and combinations of 600uM TMZ with 25uM and 50uM of AZD5582 for treated explants, and an equal volume of DMSO and water for controls. GB-EXPs in matrigel were exposed for 72hr for NAD(P)H -FLIM experiments.

#### Dose response assay

Cell viability assays were performed using the RealTime-Glo™ MT Cell Viability Assay according to the manufacturer’s protocol (Promega, Madison, WI, USA). Cells were seeded in standard media in white-walled 96-well plates at a density of 5000 cells/well and were allowed to attach overnight. The day after cells were treated with 16 combinations of TMZ and AZD5582, obtained combining 0, 10, 100 and 500 uM of TMZ and 0, 2, 5, 50 uM of AZD5582. Cells were also incubated with RealTime-Glo™ MT Cell Viability reagent. Luminescence was then measured at 24, 48, and 72 h after treatment using a multiwell plate reader (Tecan, Mannedorf, Switzerland). Data were normalized using combination of TMZ-AZD5582 0-0 uM for each time point and then analyzed using SynergyFinder online tool.

## DATA ANALYSES

### RNASeq transcriptome analysis

All analyses were conducted using the R software, version 4.2.0, within the RStudio 2022.07.02+576 suite.

#### Alignment and Annotation

FastQ files obtained from the NextSeq500 run were analysed for quality using the FastQC software (http://www.bioinformatics.babraham.ac.uk/projects/fastqc/). Reads were mapped against the reference genome (Hg19) by using STAR aligner (version 2.5.3a). Read counts on known human genes were calculated by feature Counts version 1.5.1.”Read counts were generated during the alignment process, and the raw count matrix was obtained using an R script.

#### Differential Expression Analysis

since the RNA-seq data of the 33 GBM samples were batch corrected, we normalized them and performed a differential expression (DE) analysis using the DESeq2 algorithm [17]. To reduce the impact of low counts genes, we corrected the estimated log-fold values using the normal shrinkage method. We identified differentially expressed genes by using a fold change value greater than 1.5 and a p value adjusted for multiple comparisons lower than 0.1.

### Public brain tumor datasets explorations

**GLIOVIS** data visualization and analysis tool for brain tumor public datasets was used for retrieving mRNA data, investigating survival curves, perform optimal cut-off searching and downloading data necessary to run survival analyses off the portal on GraphPad Prism 9.5.1 software [18]. The data processing for mRNA analyses consisted in: 1) Microarray. For Affymetrix expression arrays (HG-U133_Plus_2, HG-U133A, HG_U95Av2 and HuGene-1_0-st) the raw CEL files (when available) were downloaded from the respective sources and the analyses were completed in R using the Bioconductor suite. The ‘affy’ package was used for robust multi-array average normalization followed by quantile normalization. For genes with several probe sets, the median of all probes had been chosen. In case the raw data were not available and for other expression array platforms (Illumina beadchip, Agilent-4502A), the available pre-processed expression matrixs were used. 2) RNA-seq. The normalized count reads from the pre-processed data (sequence allignment and transcript abundance estimation) were log2 transformed after adding a 0.5 pseudocount (to avoid infinite value upon log transformation). Kaplan–Meier survival Analysis are performed using the ‘survival’ package in R. Hazard ratios were determined utilizing the coxph function from the ‘survival’ package. To generate HR plot we partially use some code previously described in *Cutoff finder*. For GBM classification: CIMP status has been determined by **Support Vector Machine (SVM)**, using TCGA GBM as training dataset. For the GBM subtype classification we use the 3 TCGA expression subtypes (Classical, Mesenchymal, Proneural) as defined in Wang et al 2017[19]. The Subtype calls generated by Support Vector Machine (SVM), are done performing a linear Kernel with 10-fold cross validation, using the ksvm function of the ‘kernlab’ package. As training dataset for the GBM we use the TCGA GBM samples described in Wang Q. et al. 2017. As training dataset for the LGG we use the TCGA LGG samples described in NEJM[20].

### GraphPad Prism 9.5.1 analyses

Survival analyses were all run on the GraphPad Prism software. A survival table is entered with information for each subject. Prism then computes percent survival at each time, and plots a Kaplan-Meier survival plot (and also compares survival with the log-rank and Gehan-Wilcoxon tests). All other summary data are presented as means ± s.d. All statistical analyses were performed in R and GraphPad Prism software (GraphPad 7.0, Software, La Jolla, CA). Sample size (n) values used for statistical analyses are provided in the figures and supplementary figures. Individual data point are graphed or can be found in Source Data. Tests for differences between two groups were performed using Student’s two-tailed unpaired t-test or log–rank test, as specified in the figure legends. No data points were excluded from the statistical analyses. Significance was set at P < 0.05 (for all other experiments). Differences were considered statistically significant when p < 0.05 and represented as: *p < 0.05, **p < 0.01 and ***p < 0.001.

### Jamovi software analyses

Uni- and multivariable Cox regression analyses were performed using the Jamovi package. The jamovi project (2022). jamovi. (Version 2.3) [Computer Software]. Retrieved from https://www.jamovi.org.

### Availability of data and material

All the data that support the findings of this study are available from the corresponding author upon request

## RESULTS

### NAD(P)H -FLIM based TMZ response assessment and RNAseq analysis in an extended GBM Italian case study

In this study, we included an additional group of 15 glioblastoma tumors to supplement the GBM population examined in our recent investigation [10]. These 15 new samples (GB19-GB33) comprised 4 primary and 11 recurrent glioblastoma tumors (Table 1). Following the same approach described in our previous study, the tumor samples were cultured and subjected to TMZ treatment for 72 hours using the live GB-EXP culturing procedure. For each patient, we TMZ treated and analyzed a minimum of 14 to a maximum of 40 GBM explants of similar dimensions ranging from 70 to 300μm, employing the NAD(P)H-FLIM metabolic imaging approach. The metabolic status of cancer cells is recognized as an early predictor of their response to treatment [11, 21]. The intracellular metabolic cofactor, NAD(P)H (reduced form of nicotinamide adenine dinucleotide), exists in protein-bound or protein-free states within the cell, and these states influence its fluorescence decay. Typically, bound NAD(P)H exhibits longer lifetimes than free NAD(P)H. In cancer research, metabolic shifts have been explored, and studies have reported an increase in NAD(P)H fluorescence lifetimes (indicating an increase in the NAD(P)H bound/free ratio) as cells become less proliferative and display drug responsiveness after treatment [21]. FLIM data was collected 72 hours post-treatment, and out of the 15 tumor samples, 7 were identified as TMZ responders (Resp) and 8 as non-responders (Non-Resp) with varying percentages of drug response (%DR) (Table 1).

To confirm previous gene expression findings [10], we conducted whole RNAseq analysis on the new 15 samples [10]. Consequently, the gene expression data analysis encompassing a total of 22 primary and 11 recurrent glioblastoma samples, consisting of 16 TMZ Resp and 17 TMZ Non-Resp cases (Table 1), revealed that several genes maintained consistent directional regulation of expression but lost statistical significance (Table S1) compared to the earlier results. We identified a total of 52 genes that were differentially expressed with statistical significance (adjusted p-value < 0.05) (Table S1). Among the top ten most statistically significant genes, we confirmed the upregulation of BIRC3, which exhibited a 5-fold increase in expression in the TMZ Non-Resp group compared to the Resp group (Figure 1A, Table S1). When the samples were subdivided into primary (n=22) and recurrent tumor samples (n=11), the association between BIRC3 over-expression and TMZ resistance remained consistent (Figure 1B). Among the glioblastoma samples tested, we had three matched pairs of primary and recurrent tumors. As depicted in Pair1 and Pair 3 of Figure S1, the expression of BIRC3 consistently increased in the transition from a TMZ Resp to a TMZ Non-Resp phenotype in all three matched pairs, as expected. The upregulation of BIRC3 in TMZ-resistant tumors was further validated at the protein level through immunohistochemistry (IHC), comparing TMZ Non-Resp (Figure 1C) to TMZ Resp tumors (Figure 1D).

**Figure 1.**
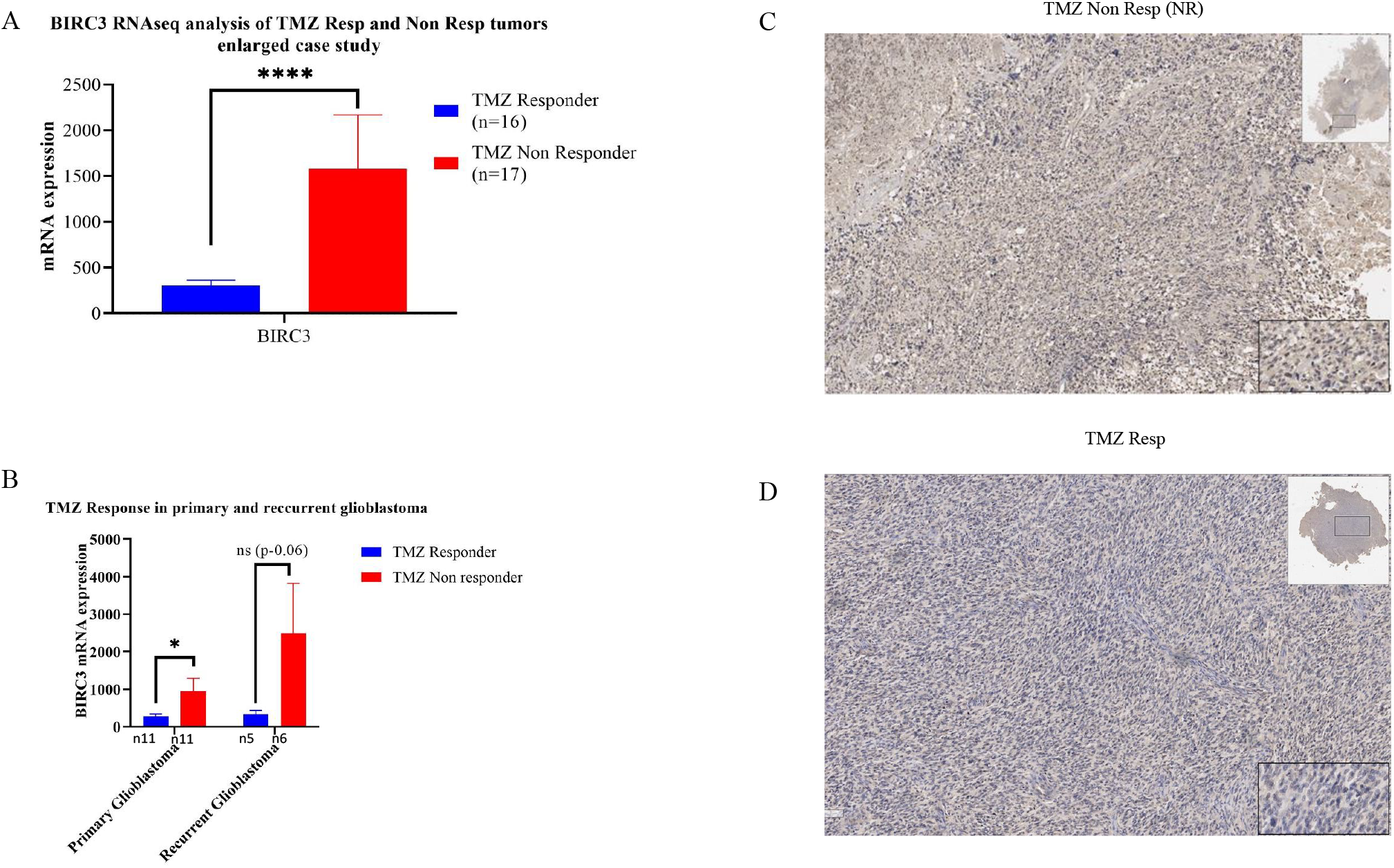
BIRC3 mRNA expression level by Whole Transcriptome Analysis. A, Statistical significant upregulation of BIRC3 mRNA (pvalue 0.00001, padj 0.02 by by Wald test)) in TMZ Non Responder tumors. B, BIRC3 mRNA levels in GBM samples stratified in primary and recurrent tumors (pvalue0.04 and 0.06 respectively by Student’s two-tailed unpaired t-test). C and D, BIRC3 protein expression patterns assessed by IHC revelead higher expression in TMZ Non-Resp tumor (A) compared to TMZ Resp tumor (B).

### Analysis of BIRC3 Expression Data in Publicly Available GBM Case Studies

To explore public brain tumors expression datasets, we exploited the GlioVis online application for data visualization and analysis [18]. As shown in Figure 2A BIRC3 expression was analyzed in several gliomas from the Rembrandt dataset [22]. Our findings reveal a statistically significant upregulation of BIRC3 expression in GBM tumors compared to lower grade gliomas and normal brain tissue, suggesting its potential involvement in tumor progression. Additionally, the IVY-GAP dataset (IVY glioblastoma atlas project, http://glioblastoma.alleninstitute.org) highlights a statistically significant association between the upregulation of BIRC3 expression and the presence of pseudopalisading cells around necrosis in different intra-tumor regions tested, as illustrated in Figure 2B. In TMZ Non-Resp cases, we observed a higher concentration of BIRC3 protein signal around necrotic areas through IHC, consistent with the findings reported in the IVY datasets (Supplementary Figure S2A). Conversely, TMZ-Resp tumors in peri-necrotic areas exhibited low or no expression of BIRC3, as depicted in Supplementary Figure S2B. Furthermore, Supplementary Figure S2 C and D demonstrate the absence or minimal expression of BIRC3 in the tumor core regions of both tumor types.

**Figure 2.**
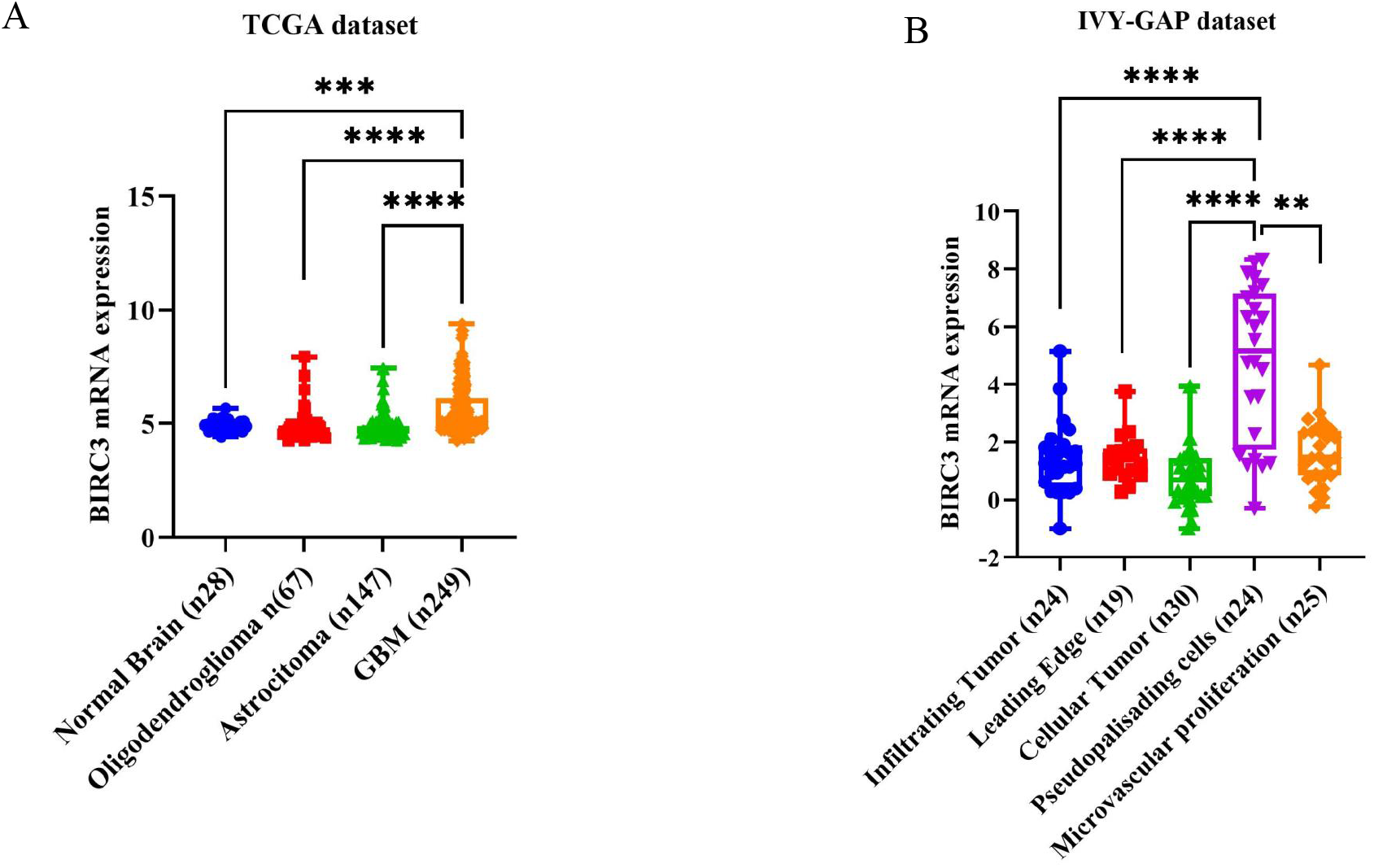
BIRC3 mRNA distribution inter and intra tumors according to GlioVIs portal. A, BIRC3 expression was analyzed in several gliomas from the Rembrandt dataset showing a statistical significant upregulation in GBM tumors compared to lower grade gliomas. B, BIRC3 expression is reported to be statistically significantly associated with pseudopalisading cellular structures compared to other intratumor regions according to IVY-GAP dataset (IVY glioblastoma atlas project, http://glioblastoma.alleninstitute.org). Pvalues were calculated by Student’s two-tailed unpaired t-test.

### Survival Analysis Based on BIRC3 mRNA Expression in Publicly Available Datasets

We examined the association between BIRC3 expression and overall survival (OS) probability in three glioblastoma datasets (TCGA) available through the same web application tool (GlioVis). These datasets consisted of GBM tumors derived from clinically characterized patients and were analyzed using three different transcriptome technical approaches: Agilent-4502A (n=488), Affymetrix HG-UG133 (n=525), and RNAseq (n=155) platforms (Table S2, S3, S4). In Figure 3A, B, and C, Kaplan-Meier analyses demonstrate that the median value of BIRC3 expression serves as a threshold to identify a higher-risk group (BIRC3 High) and a lower-risk group (BIRC3 Low) in terms of OS, with patients in the high BIRC3 expression group exhibiting a worse prognosis compared to those with lower BIRC3 levels, which were associated with longer survival. The median OS values between the curves were statistically significantly different for the Agilent-4502A and Affymetrix HG-UG133 datasets but not for the RNAseq dataset, as depicted in Figure 3A, B, and C. To determine if a different threshold could provide more informative OS probability curves, we utilized the *optimal-cut-off* survival analysis tool available within the same GlioVis portal [18] (http://gliovis.bioinfo.cnio.es/). Figure 3D, E, and F demonstrate that by selecting a different threshold, we could identify a more precise population of patients with better or worse OS probabilities. The population characterized by BIRC3 expression levels below the *optimal-cut-off* exhibited longer OS, which statistically significantly deviated from the population with the worst prognosis in all three datasets, compared to the survival analyses based on the BIRC3 expression median value. In the TCGA large case studies we assessed also the complementarity of BIRC3 expression levels and the methylation status of the MGMT promoter. Supplementary Figure S3A, B, and C illustrate that the MGMT promoter methylation status alone was a statistically significant predictor of outcome in all three TCGA datasets. Survival analysis performed using a combination of BIRC3 Low and High scores (optimal-cut-off approach) with MGMT methylation status revealed a synergistic outcome prediction in all three datasets (Figure S3E, F, G). The combination of these two molecular parameters identified four groups of patients, with the BIRC3 Low and methylated MGMT group displaying the longest median overall survival, which was statistically different from the group of patients with BIRC3 High and unmethylated MGMT in all three datasets (Figure S3E, F, G). The median OS changed from 17.6 to 22.7, 17 to 21.1, and 15.9 to 25.4 months in the Agilent-4502A, Affymetrix HG-UG133, and RNAseq TCGA datasets, respectively (Figure S3E, F, G), when comparing the MGMT methylated status alone or in combination with the BIRC3 Low score. The prognostic significance of BIRC3 expression was further validated through univariate and multivariate analysis (Figure S4), demonstrating that BIRC3 expression plays an independent prognostic role in glioblastoma. To further validate the prognostic role of BIRC3 expression, we analyzed its significance in a separate dataset, namely the Chinese Glioma Genome Atlas (CCGA), which was analyzed using the RNAseq approach and accessible through the GlioVis portal (Table S5). Supplementary Figure S5A illustrates that BIRC3 is highly expressed in glioblastoma (GB) compared to other histologies, confirming its involvement in tumor progression, consistent with the findings from the TCGA dataset. The median overall survival (OS) of the CCGA patients was 15.9 months (data not shown). Furthermore, survival analysis once again demonstrated a statistically significant association between BIRC3 expression and survival, showing the same expression pattern observed in the TCGA datasets. The OS median for patients with low BIRC3 expression was 19.8 months, while those with high BIRC3 expression had a median OS of 13.9 months (Figure S5B).

**Figure 3.**
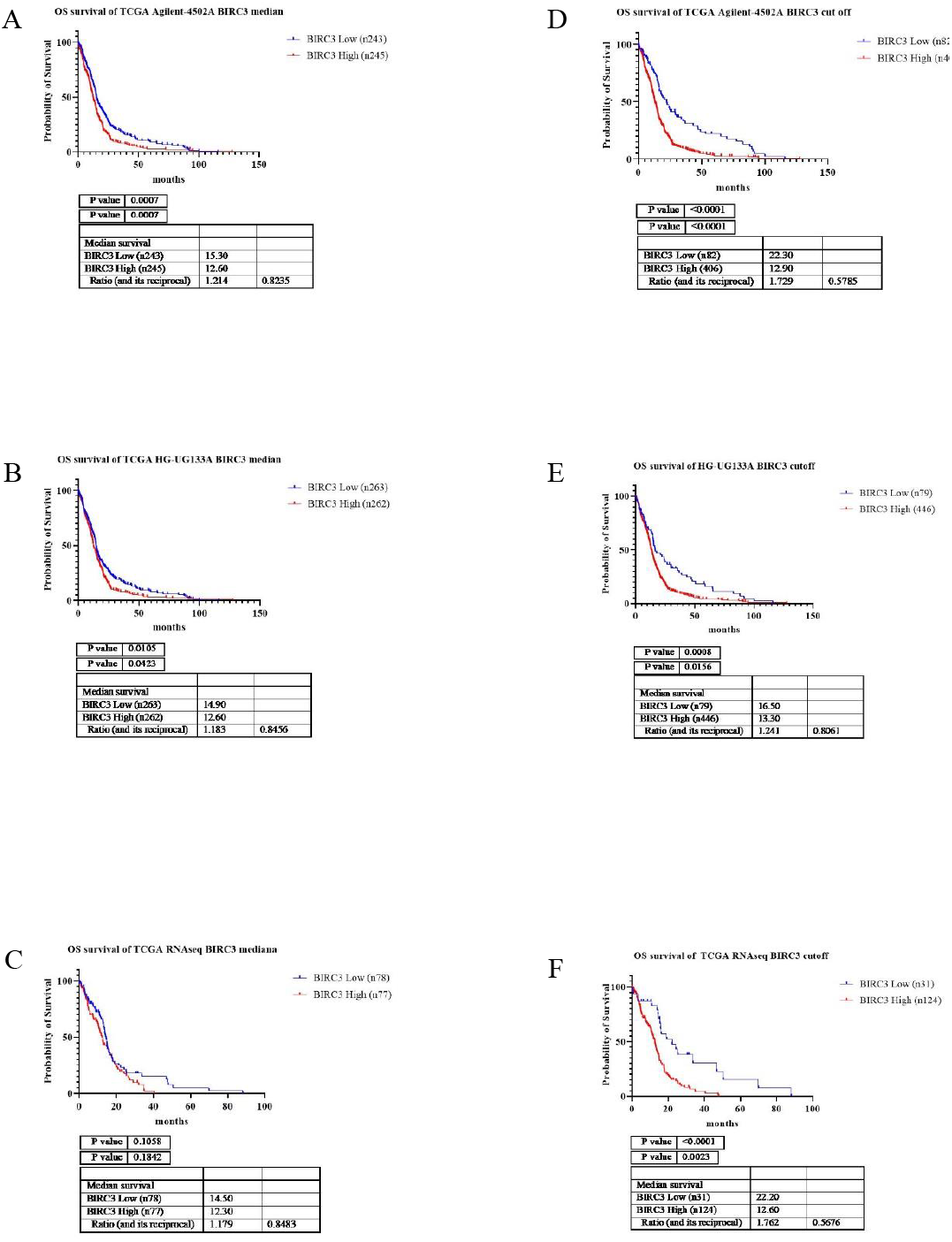
Kaplan Meier survival analyses based on transcriptome data of TCGA datasets obtained with three different technical approaches, Agilent-4502A (n488), Affymetrix HG-UG133 (n525) and RNAseq (n155). A-C, Survival analyses performed utilizing the median BIRC3 expression value as a threshold for discriminating a High and Low expression group. D-E, Survival analyses performed exploiting the *optimal*-*cut-off* tool provided by the GlioVis portal. For each analysis two pvalues are calculated with Logrank/Mantel-Cox and Gehan-Breslow-Wilcoxon, provided by GraphPad software.

### Association between BIRC3 mRNA Expression and Treatment Response in Patients from the TCGA RNAseq Dataset

We utilized the RNAseq dataset from TCGA to investigate the relationship between BIRC3 expression and treatment outcomes in glioblastoma multiforme (GBM). The dataset comprised 142 patients, categorized as follows: 95 underwent chemo and radiotherapy (Stupp protocol), 19 received radiotherapy alone, 2 received chemotherapy alone, and 26 received no treatment at all (Table S4). [23]. Through an optimal-cut-off approach, we identified that patients with a low BIRC3 score exhibited a favorable response to the Stupp protocol, confirming its association with a good prognosis (Mantel-Cox p=0.1/Gehan-Breslow-Wilcoxon p=0.04) (Figure 4a,b). Moreover, within the same group of GBM patients, BIRC3 expression showed a strong association (Mantel-Cox p=0.01/Gehan-Breslow-Wilcoxon p=0.03) with progression-free survival (PFS) (Figure 4c,d). Survival analysis of the untreated patient group further demonstrated that tumors with lower BIRC3 expression (optimal-cut-off approach) correlated with longer survival, irrespective of treatment, suggesting that BIRC3 expression reflects both a tumor’s genetic background and its intrinsic aggressiveness, as well as predisposition to TMZ treatment resistance (Figure 4e,f,g,h). These findings collectively indicate that the BIRC3 expression levels can serve as a predictive model for prognosis and therapeutic response in GBM patients.

**Figure 4.**
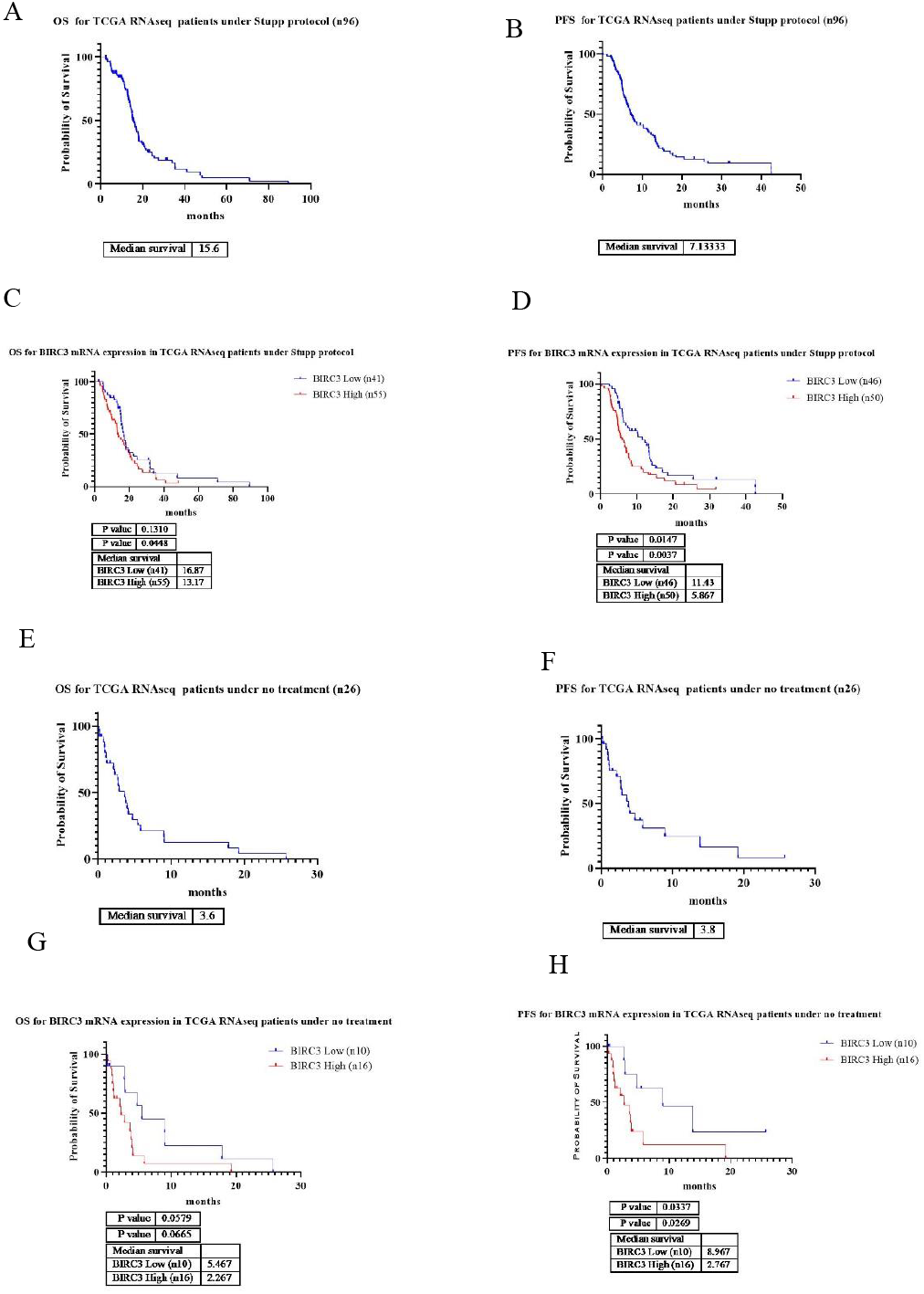
BIRC3 expression and treatment response in patients of TCGA RNAseq dataset. A and B, Overal survival (OS) and Progression free survival (PFS) in a selection of patients that were all admistered the Stupp protocol. C and D, Kaplan Meier survival analyses based on BIRC3 mRNA expression (optimal cut-off) in the patients undergone Stupp protocol showing an association of lower levels of BIRC3 with better OS and PFS and higher levels of BIRC3 mRNA expression with worse prognosis. E and F, Overal survival (OS) and Progression free survival (PFS) in a selection of patients that did not received any treatment. G and H, Kaplan Meier survival analyses based on BIRC3 mRNA expression (optimal cut-off) in not treated patients showing a significant statistical association between levels of BIRC3 mRNA expression and OS or PFS. For each analysis two pvalues are calculated with Logrank/Mantel-Cox and Gehan-Breslow-Wilcoxon, provided by GraphPad software.

### Prognostic Validation of BIRC3 Expression in European Private GBM Case Studies

The findings on BIRC3 obtained from publicly available datasets were further validated through the analysis of a private French case study (n=229) (Table S6) using RNAseq, and an Italian case study (n=65) (Table S7) using Real-Time PCR. The French case study consisted of IDH1/2 wildtype primary glioblastoma. Among the 228 French GBM tumors, MGMT promoter methylation status was available for 113 tumors, with 66 tumors showing MGMT methylation and 47 tumors being unmethylated (Table S6). The Italian case study included 47 primary and 18 recurrent IDH1/2 wildtype glioblastoma tumors, and methylation status information was not available (Table S7). All patients in both case studies underwent the Stupp protocol. Survival analysis of the 228 French glioblastoma tumors revealed an overall survival (OS) of 18.7 months and a progression-free survival (PFS) of 11.3 months (Figure 5A,B). When exploring the prognostic role of BIRC3 expression, lower levels of BIRC3 expression (median expression threshold) were significantly associated with better prognosis for both OS and PFS, as depicted in Figure 5C,D. The OS and PFS improved to 20.8 and 12.6 months, respectively, compared to the general survival assessment, as observed in the publicly available TCGA datasets. MGMT methylation status alone exhibited a stronger prognostic role compared to BIRC3 status (Figure 5E,F), but the combination of both molecular parameters (Figure 5G,H), as observed in the publicly available TCGA datasets, provided more informative survival profiling, particularly for the median probability of PFS, which increased from 15 months when calculated using MGMT methylation status alone to 19.6 months when combined with BIRC3 status (Figure 5F,H). Furthermore, MGMT methylation status alone predicted an OS of up to 65 months in 10% of patients (Figure 7E, arrow), while the combination with a BIRC3 low score increased the percentage to 20%, as shown in Figure 5G (arrow). Univariate analysis of the French case study demonstrated that BIRC3 was a significant prognostic factor associated with OS and PFS survival, independent of age or sex, but dependent on the MGMT methylation status (data not shown).

**Figure 5.**
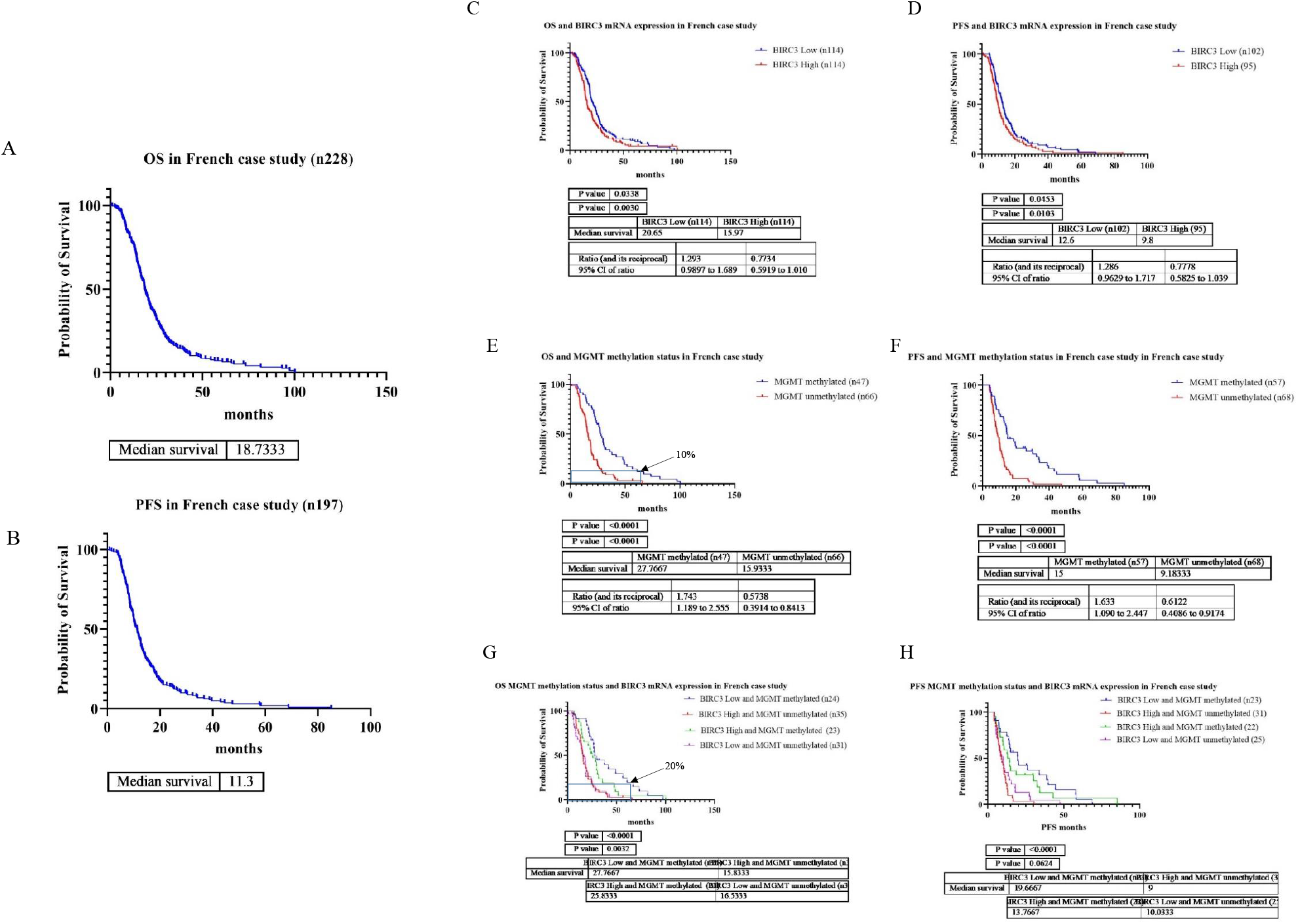
BIRC3 mRNA expression based survival data analysis in private French GBM case study. A and B, Overal survival (OS) and Progression free survival (PFS). C and D, Kaplan Meier survival analyses based on BIRC3 mRNA expression (median) improves prognosis, showing an association of lower levels of BIRC3 with better OS and PFS and higher levels of BIRC3 mRNA expression with worse prognosis. E and F, Kaplan Meier survival analyses based on MGMT promoter methylation status) improves prognosis, showing that MGMT methylated tumors are associated with a better OS and PFS. G and H, Kaplan Meier survival analyses based on the combination of MGMT methylation status and BIRC3 expression levels. Results show that the combination of the two parameters stratifies patients in different survival groups with an increase in percentage of patients with longest survival when both MGMT meth and Low BIRC3 are combined. For each analysis two pvalues are calculated with Logrank/Mantel-Cox and Gehan-Breslow-Wilcoxon, provided by GraphPad software.

The OS and PFS in the Italian case study were 15.7 and 9.4 months, respectively (Figure 6A,B). Real-time PCR analysis of BIRC3 expression was performed on 65 Italian GBM samples (Table S7). OS and PFS data were available for 55 samples, and survival analysis confirmed that BIRC3 expression was statistically significantly associated with survival in glioblastoma patients (Figure 6C,D). Consistent with expectations, the OS and PFS probability curves in the Italian case study, when divided based on the median expression value of BIRC3, exhibited statistically significant differences in median survival. The group with lower BIRC3 expression had a median OS and PFS of 21.9 and 11.4 months, respectively (Figure 6C,D), while the group with higher BIRC3 expression showed worse prognosis, with median OS and PFS of 14 and 8 months, respectively (Figure 6C,D). The same statistically significant association between BIRC3 expression and OS or PFS, with the same directionality, was observed when analyzing only the primary tumors from the Italian case study, as shown in Figure S6a,b.

**Figure 6.**
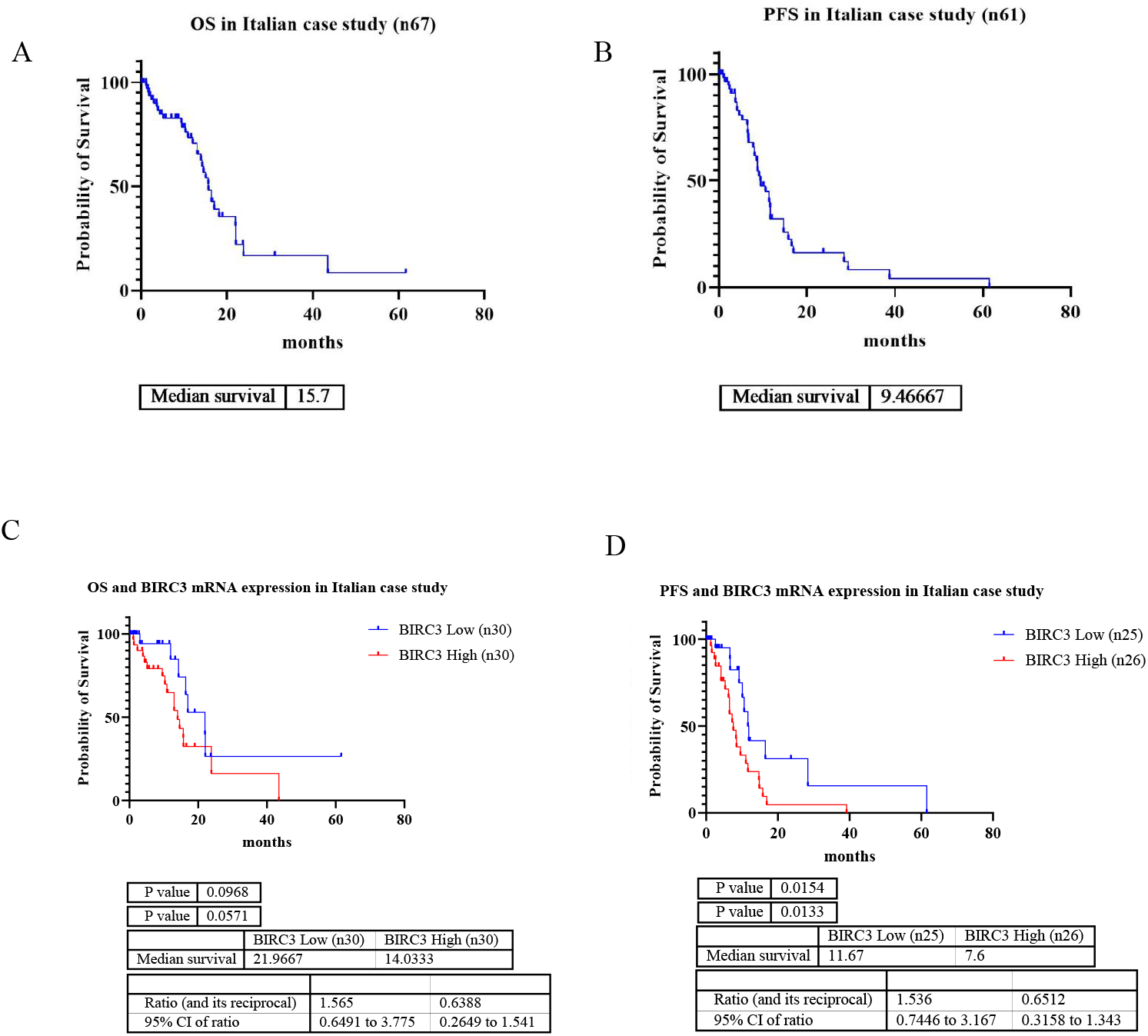
BIRC3 mRNA expression based survival data analysis in private Italian GBM case study. A and B, Overal survival (OS) and Progression free survival (PFS). C and D, Kaplan Meier survival analyses based on BIRC3 mRNA expression (median) improves prognosis, showing an association of lower levels of BIRC3 with better OS and PFS and higher levels of BIRC3 mRNA expression with worse prognosis. For each analysis two pvalues are calculated with Logrank/Mantel-Cox and Gehan-Breslow-Wilcoxon, provided by GraphPad software.

### Reversal of TMZ-Resistant Phenotype in T98G GBM Commercial Cell Lines and Patient-Derived Live Tumors through Targeting BIRC3 with AZD5582

#### GBM commercial cell lines

T98G are renown TMZ resistant GBM commercial cell lines [9]. Western blot analysis revealed robust expression of the BIRC3 protein in T98G cells (Figure 7A), and immunohistochemistry showed predominant cytoplasmic localization (Figure 7B). AZD5582, a potent antagonist of apoptosis inhibitors specifically targeting BIRC2 and BIRC3, was developed by AstraZeneca in 2013 [15]. To overcome TMZ resistance, we conducted dose-response and drug synergy assays on T98G cells using TMZ, AZD5582, and their combination. Figure 7C and 7D depict the inhibition curves of TMZ and AZD5582-treated T98G cells at doses of 0μM, 10μM, 100μM, and 500μM, and 0μM, 2μM, 5μM, and 50μM, respectively, at 72 hours. As expected, T98G cells exhibited resistance to TMZ, with only 60.17% proliferation inhibition observed at the highest dose of 500μM (Figure 7C,E). In contrast, AZD5582 demonstrated significant inhibition of T98G proliferation, reaching 61.16% at 2μM and 99.6% and 99.8% at 5μM and 50μM, respectively (Figure 7D,E). The lack of proliferation inhibition observed at TMZ doses of 10μM and 100μM was reversed, resulting in 27.19% and 39.45% inhibition, respectively, when combined with 2μM of AZD5582 (Figure 7E). The loss of TMZ resistance was further enhanced when 5μM of AZD5582 was added to the assay (Figure 7E). A synergy map assay can be extrapolated from the dose-response assay results, highlighting the antagonist effect of the combination of 2uM AZD5582 and any TMZ doses (Sinergy score ranges from -24,58 to -30.8) (Figure 7F). However, the antagonistic effect observed at 2μM AZD5582 changed to an additive effect when 5μM of AZD5582 was combined with 100μM and 500μM of TMZ (synergy score of -4.5 and -4.94, respectively) (Figure 7F). These results demonstrate that AZD5582 treatment efficiently overcomes TMZ resistance in T98G GBM cell lines when combined with TMZ treatment through the antagonistic effect of the two drugs. Using an apoptosis detection assay (Beckman Coulter), we assessed the presence of early and late apoptotic cells after 72 hours of treatment with TMZ (100μM), AZD5582 (2μM), or a combination of both (Figure 7G). As shown in Figure 7G, only the combination treatment exhibited a statistically significant induction of apoptosis compared to controls and single drug treatments, indicating the antagonistic effect of AZD5582 [15, 24] in overcoming apoptotic inhibition mediated by BIRC3. Additionally, it is worth noting that cellular BIRC3 expression was previously described by Wang et al. in 2016 to be progressively induced by TMZ exposure during the onset of drug resistance.

**Figure 7.**
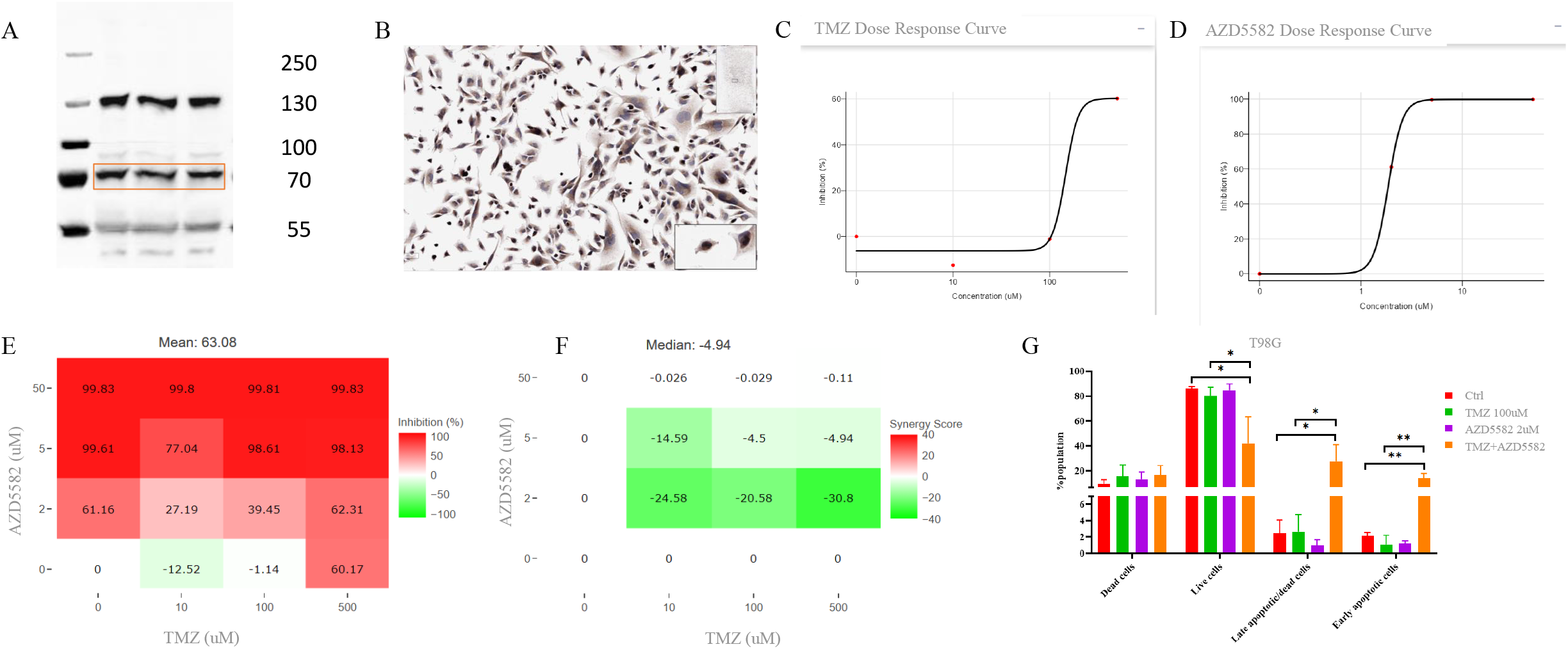
AZD5582 reverts TMZ resistant phenotype in T98G GB commercial cells. A, Western blot of BIRC3 protein expression in T98G cells (performed in triplicate). B, BIRC3 protein expression in T98G cells in culture by immunocytochemistry (ICC). ICC shows a cytoplasmic and nuclear localization. C and D Dose response curves of TMZ and AZD5582 administered alone to T98G cells. E, Proliferation inhibition map of TMZ and AZD5582 treatments alone and in combination at different doses. F, Synergy map TMZ and AZD5582 treatments alone and in combination at different doses. Synergy scores are representative of a antagonist effect when <-10. Additive effect between >-10-<10 and synergistic effect <10. Dose response curves, inhibiton and synergy maps were obtained with Synergy Finder software G, Flow cytometry Annexin based apoptotic detection (Beckman Coulter) on T98G cells treated with TMZ and AZD5582 at 72hr show a statistical significant increase in apoptosis (early and late apoptotic cells) after combining TMZ with anti BIRC3 AZD5582.

#### GBM patients-derived tumors

Among the patients-derived live tumors, four GBM samples were selected, all characterized as TMZ non-responsive (NR) based on NAD(P)H-FLIM drug response assessment (Figure 8A). As depicted in Figure 10a, three samples (GB22, GB29, and GB76) exhibited a 20-fold higher expression of BIRC3 compared to GB37, which, despite its low BIRC3 expression, also displayed TMZ non-responsiveness. These four patient-derived tumors were cultured as GB-EXPs [10] and treated for 72hr with 600uM TMZ, 25uM and 50uM AZD5582 singularly. At the same time, we tested the combination of 600uM TMZ with 25uM and 50uM of AZD5582. Higher doses of AZD5582 were used considering the 3D structure of the GB-EXPs and penetration limitations due to matrigel embedding, as observed in in vitro TMZ assays [25]. The response to each treatment was evaluated using NAD(P)H-FLIM metabolic imaging following the procedure outlined in our previous study [10]. All samples were confirmed to have a 0%DR when treated with 600uM of TMZ as shown in Figure 10b. A concentration of 25μM AZD5582 was highly effective (%DR HR) for three tumor samples, except for GB37 (0% DR, NR), which exhibited a very low expression of the BIRC3 target specific to AZD5582, compared to the other three samples (Figure 8A). A higher concentration of 50μM AZD5582, which was cytotoxic in 2D cell lines (Figure 7E), resulted in measurable %DR in all samples, with the lowest response observed in GB37 (34% DR, MR), and the highest in sample GB22 (90% DR, HR) (Figure 8B). The combination of 25μM AZD5582 and 600μM TMZ successfully reversed the resistant phenotype in the three TMZ NR GB samples with high BIRC3 expression (Figure 8B), while no change was observed in GB37 with low BIRC3 expression. The TMZ NR phenotype of GB29 and GB76 was converted to a TMZ HR phenotype, while GB22 acquired a LR phenotype (19%DR). Similar results were obtained when the combination included the higher dose of 50μM AZD5582 with 600μM TMZ. In this case, the lowest %DR was observed in GB37 (12% DR, LR), suggesting that BIRC3 expression plays a determinant role in the action of AZD5582. As observed in 2D cell lines (Figure 7F), these results demonstrate that the combination treatment of AZD5582 and TMZ efficiently overcomes TMZ resistance in three patient-derived GB-EXPs with high BIRC3 expression through an antagonistic effect of the two drugs, as evidenced by the lack of a consistent improvement in %DR compared to AZD5582 treatment alone (Figure 8B). Moreover, the differential drug response behavior between the three samples (GB22, GB29, GB76) and GB37 suggests that BIRC3 expression levels could potentially be used to selectively stratify TMZ+AZD5582-responsive tumors among TMZ NR GBM tumors. Representative bright-field and FLIM images, along with corresponding NAD(P)H free/bound distribution curves [10] of controls and treated GB-EXPs are shown in Figure 8C-J for sample GB37 and sample GB29 that resulted TMZ+AZD5582 NR and HR respectively, at 72hours post treatment.

**Figure 8.**
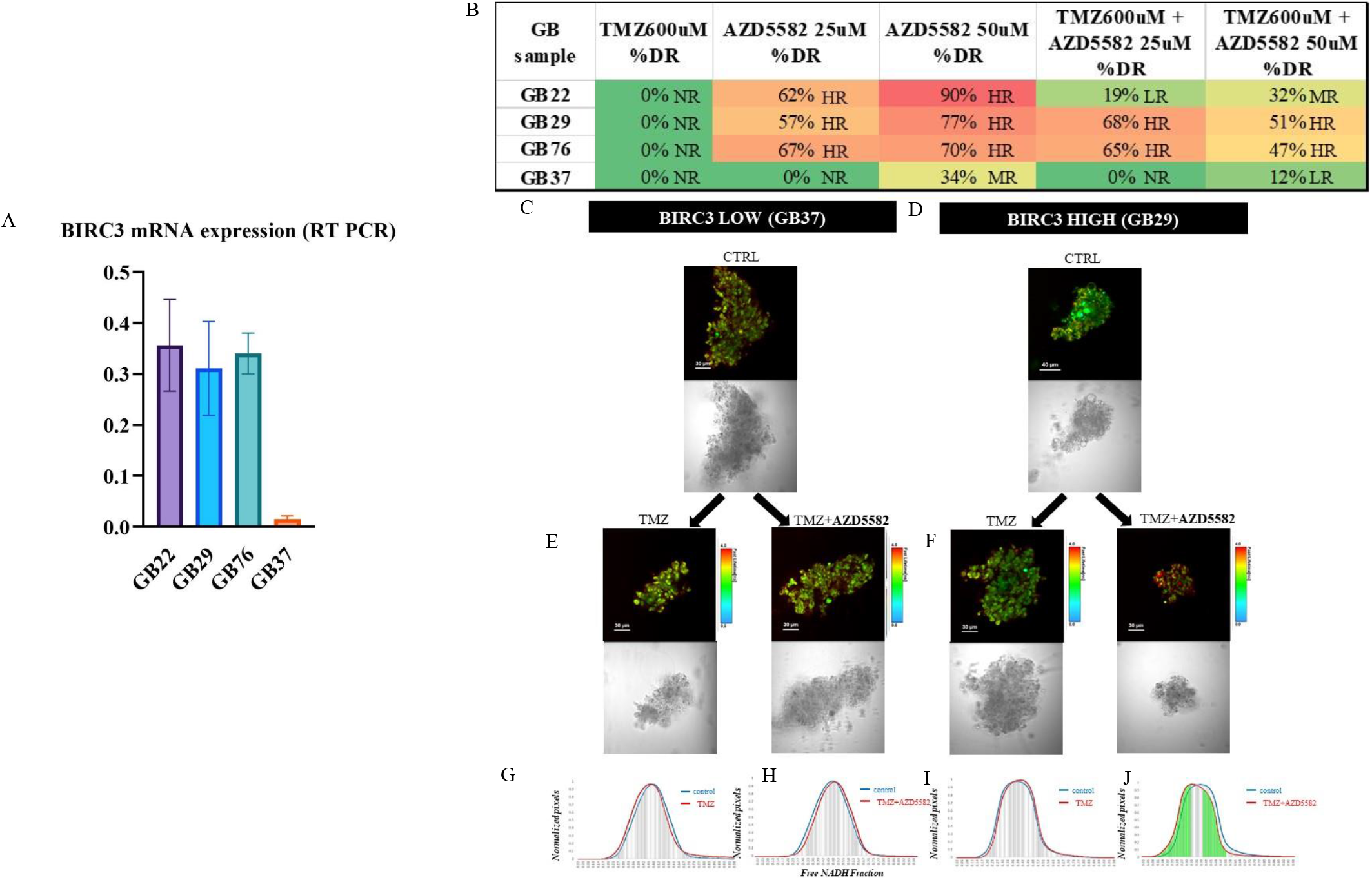
AZD5582 reverts TMZ resistant phenotype in patients-derived GBM live explants (GB-EXPs) in culture. A, BIRC3 mRNA expression levels (RT-PCR) in 4 GBM tumors (RT-PCR done in triplicate). B, Percentage of Drug Response (%DR) in GB-EXPs measured with NADH-FLIM metabolic imaging at 72hr post treatment for TMZ and AZD5582 administered alone and in combination (an average of 18 EXPs per tumor sample per treatment were assessed by NADH-FLIM approach). C, D Representative images of GB37explant with low expression of BIRC3 (C) and GB29 (D) with higher levels of BIRC3 mRNA expression. C,D FLIM and bright-field representative images of non-treated GB-EXPs samples. E and F, FLIM and bright-field representative images of treated GB37 (E) and GB29 (F) with TMZ and combination of TMZ and AZD5582. G-J, TMZ treatment showed no response when administered alone in both GB37 and GB29 samples as shown by a complete overlap of NADH free/bound distributions (G, I). The combination of AZD5582 with TMZ reverts the resistant phenotype of GB29 with high levels of BIRC3 AZD5582 specific target, shown by a significant statistical shifts of the NADH free/bound distribution curve of the treated samples compared to controls (J) while no change in TMZ resistance was observed in GB37 (H) characterized by a 20-fold lower level of BIRC3 mRNA expression (A).

## 2) DISCUSSION

Glioblastoma (GBM) is a highly aggressive cancer with poor prognosis that requires improved treatment options. There is a lack of reliable biomarkers to predict treatment response and prognosis. While several molecular markers have been proposed as potential biomarkers for GBM, their implementation into clinical settings is slow and impeded by tumor heterogeneity [26]. Standard GBM treatment involves surgical resection followed by radiation and concomitant temozolomide (TMZ), then adjuvant TMZ to target remaining tumour cells [23]. Recurrence is, on average, observed 7–10 months, post treatment [27]. The median overall survival (OS) of GBM patients is 12–14 months, largely due to the cancer’s invasiveness and resistance to treatments, even after treatment with TMZ and radiation [26, 27]. To date, few molecular biomarkers have been discovered. These include O6 -methylguanine DNA methyltransferase (MGMT) promoter methylation [28], Isocitrate dehydrogenase 1 (IDH1) mutation[29], mutations in the promoter region of the telomerase reverse transcriptase (TERT) gene [30], and amplification and/or overexpression of EGFR [31]. These potential prognostic molecular biomarkers have value for GBM patient management or are informative in the context of standard of care TMZ and radiation treatment [28]. However, for a subset of patients, the outcome is not well predicted by these markers, and may be better or worse than expected. This highlights the need to investigate other biomarkers associated with prognosis and response to treatment[26]. Even with the addition of new therapies, TMZ has become a cornerstone of GBM treatment, but it is unfortunately also a key factor in tumor resistance and recurrence. Understanding and combating TMZ resistance is further complicated by the fact that resistance can either be inherent or acquired after initial treatment [32-34]. In this study we used NAD(P)H -FLIM metabolic imaging to stratify patient-derived tumors into responders and non-responders to TMZ treatment (Table.1). FLIM measures NAD(P)H lifetimes, representing a powerful non-invasive tool to monitor, in real time, metabolic activities in living cells and tissues [6]. Usually, the stratification of tumor response to a therapy is carried out on the basis of clinical results, thus introducing several confounding factors which can mask the real genetic structure of the original tumor and therefore lose more precise information. The approach we used [10], is instead based on the response to the TMZ treatment measured ex vivo directly on the patient’s tumor and therefore based only on its innate biology. This allowed the identification of several statistically significant genes differentially expressed between TMZ responsive and non-responsive tumors, which we were able to confirm even in a larger series. The BIRC3 gene encoding an apoptosis inhibitory protein, member of the IAP family proteins [35] was found to be among the top ten most significantly differentially expressed genes with a higher overexpression in our TMZ-resistant tumors at the mRNA (Figure 1A) and protein level (Figure 1C) compared to TMZ sensitive tumors. TMZ exerts its action by introducing unrepairable mutations that induce the formation of single- and double-stranded DNA breaks resulting in cell cycle arrest at G2/M and finally apoptosis [25]. Although there can be many causes behind the TMZ acquired drug resistance, a major mechanism of resistance lies in the dysfunctional survival signaling in cancer cells and the reluctance of these cells to undergo apoptosis. Indeed, the propensity of cancerous cells to avoid undergoing apoptosis has long been recognized as one of the hallmarks of cancer [36] . In a preclinical model of a human GBM cell line, Wang et al found an increase in BIRC3 expression in response to irradiation (RT) and temozolomide (TMZ) treatment [14]. In our study, we explored the mRNA expression of BIRC3 in several case studies of the TCGA and CCGA transcriptome datasets exploiting the GlioVis web application for data visualization and analysis [18]. Moreover, in the TCGA dataset, elevated BIRC3 gene expression was identified as a superior and selective biomarker of mesenchymal GBM versus neural, proneural and classical subtypes [37] and specifically, it was reported as associated with TMZ resistance, patients’ survival and stemness reprogramming in GBM cell lines [38] [14]. Our investigations show that BIRC3 progressively increases its expression from control, low grade to high grade gliomas (Figure 2a, Figure S5A) and that is specifically overexpressed in the tumor perinecrosis areas (Fig.4B and Figure S2) composed of hypoxic pseudopalisades cells with mesenchymal phenotype [39, 40], suggesting that BIRC3 represents a potential candidate marker of GBM progression and aggressiveness. Moreover, in three different transcriptome TCGA datasets, obtained with different methodological approaches we performed several Kaplan Meier survival analyses and found that lower level of BIRC3 expression are always associated significantly with a better prognosis (Figure 3A,B,C) as, also, described in a 2016 study [14]. Multivariate analysis proved that BIRC3 is an independent prognostic factor and can be proposed as a new marker of survival (Fig.S4). Our results were also obtained in the Chinese GBM dataset (CCGA) as shown in Fig.S5 supporting the cross-sectional BIRC3 prognostic role that goes beyond genetic differences of ethnicity. In patients given no treatment, low BIRC3 expression was prognostic of longer survival, indicating that its expression level may reflect a more or less aggressive tumor state as early as start (Fig. 4E-H). Taken together, these data demonstrate that BIRC3 expression, in GBM patients, can not only be predictive of therapy response but also prognostic of better outcome regardless of treatment. Therefore, the TCGA data led us to confirm BIRC3 prognostic role also in the two clinically characterized European datasets that we collected, composed of a French case study of 228 tumor samples and 65 Italian GBM tumor samples (Figure 5,6).

What sets our work apart from Wang et al. in 2016 and 2017 [14, 37] is the comprehensive investigation of the role of BIRC3 in an additional case study from the TCGA dataset, as well as two completely separate private GBM case studies collected in Europe with distinct clinical characterizations. The utilization of diverse techniques to obtain our data further enhances the validity and robustness of our results. Furthermore, the signal of BIRC3 upregulation associated with TMZ resistance emerges initially from a functional experiment done on viable tumor tissues derived directly from patients exposed to TMZ treatment. Remarkably, BIRC3 emerged as one of the most significant differentially expressed genes consistently observed throughout the entire transcriptome analysis.

Cells can undergo apoptosis in response to a variety of stimuli, such as chemotherapy or radiation-induced cellular stresses. Specifically, the final intracellular consequence of TMZ administration is inducing the cells on receiving a pro-apoptotic signal, and a variety of intracellular signaling process occur, eventually culminating in the activation of apoptosis through cysteine-dependent aspartyl specific proteases (caspases) that are responsible for effecting cell death. A high level of BIRC3 interferes with this process and counteract TMZ effect by developing resistance [14]. Thus, agents that can restore sensitivity of these cells to various pro-apoptotic cues might conceivably be employed as cancer therapies, either alone or in combination with other drugs [41]. Given the well-documented role of the IAP family of proteins in the suppression of apoptosis in cancer cells as well as the potential for a Smac mimetic to be an effective treatment for human cancers, in 2013 Hennessy et al [15] developed a novel compound inhibitor of the IAPs, the AZD5582, proven to inhibit proliferation in a subset of cancer cell lines. In the same study authors demonstrated effects of IAP inhibition in vivo and the ability of AZD5582 to exert antitumor activity in a MDA-MB-231 xenograft model [15]. Recently in 2022, the drug AZD5582 was described to selectively kill Fanconi Anemia (FA) head-and-neck squamous cell carcinomas transformed cells that overexpressed BIRC2 or BIRC3 [24]. To the best of our knowledge, based on these results, we decided to test the AZD5582 compound for the first time on T98G glioblastoma cell lines, resistant to TMZ treatment and significantly overexpressing BIRC3 compared to U87 and U118 TMZ responsive cells (Figure S7) in the attempt to revert TMZ resistance. As shown in Figure 7E,F we observed that the combination of the two drugs at different doses was able to revert the T98G TMZ resistant phenotype, due to an antagonistic effect. The combination of AZD5582 with TMZ causes a statistically significant increase in T98G proliferation inhibition (Figure 7E,F) and in apoptosis (Figure 7G) compare to TMZ when administered singularly. However, at the same time AZD5582 proliferation inhibition is reduced by the presence of TMZ as shown in Figure 7E,F. These results may explain a contrasting action which sees on one hand the inefficient action of TMZ which cannot exercise its apoptosis induction activity [9] due to the presence of a high expression of the BIRC3 inhibition which justifies the resistance of T98G to the drug itself [14]. On the other hand, the action of AZD5582 suppresses the action of BIRC3 and contrasts the TMZ resistance. The same drug response investigation was performed in patients-derived live GB-EPXs utilizing NAD(P)H - FLIM metabolic imaging approach [10]. We chose only TMZ resistant patients derived tumors, previously assessed for TMZ response. Three of the 4 tumors were expressing high levels of mRNA BIRC3 compared to the exceptional TMZ resistant case that had a 20-fold lower expression of BIRC3 (Figure 8A). We intentionally chose one of the few tumors that had a low expression of BIRC3 in contrast to the TMZ resistance association observed in this study. In GB-EPXs samples we obtained the same results observed for T98G cell lines, as TMZ resistance was reverted after combining AZD5582 to TMZ. However in the sample lacking BIRC3 expression, we had no change in TMZ resistant, suggesting that the low expression of the AZD5582 specific target prevented the drug from exerting its effect and indicating that TMZ resistance in this tumor was potentially dependent of other molecular factors not involving BIRC3 (Figure 8B,C).

Traditionally, precision oncology has relied on static features of the tumor to determine appropriate therapies. These features may include the expression of key targets or genomic analysis of mutations to identify therapeutically targetable “drivers.” However, in glioblastoma, only a small proportion of individuals have benefited from this static approach. As a result, functional precision medicine can offer valuable insights into tumor vulnerabilities beyond what is provided by static features. Our functional precision medicine approach for tumor drug assessment enables the stratification of tumors based solely on their biology [6, 10]. This results in greater sample homogeneity and the identification of more reliable and robust differentially expressed genes that can be utilized to refine drug discovery and enable personalized targeted clinical management. Indeed, on the basis of our results, BIRC3 as an inhibitor of apoptosis, represents a potential biomarker of prognosis and a predictor of TMZ treatment response in GBM, and complements MGMT status as an outcome predictor. Additional experiments are needed on larger GBM tumor case studies to establish a BIRC3 mRNA expression threshold that optimally stratifies GBM patients into low or high expression groups with probability scores for better or worse survival, respectively. Moreover, BIRC3 represents also a new therapeutic target specific of the AZD5582 drug in glioblastoma tumors resistant to TMZ treatment. The level of BIRC3 expression could be used to stratify patients who will benefit from a combination of TMZ and AZD5582. Our study provides potentially valuable leads for future clinical trials that will determine whether NAD(P)H - FLIM as a functional precision medicine approach combined with BIRC3 expression assessment can improve patients’ management and become a standard tool in glioblastoma clinical oncology.

## Acknowledgements

we thank all the patients that gave their informed consent

