## Supplementary Figures for "BIRC3: A Prognostic Predictor and Novel Therapeutic Target in TMZ-Resistant Glioblastoma Tumors"

### Figure S1

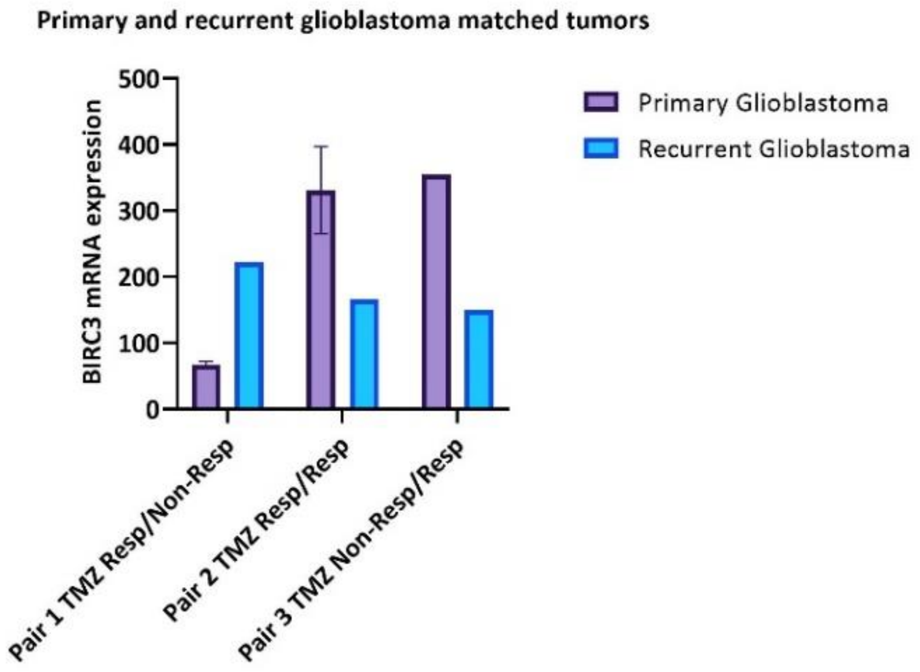

Primary and recurrent glioblastoma matched tumors
BIRC3 mRNA expression
500
400
300
200
100
0
Primary Glioblastoma
Recurrent Glioblastoma
Pair 1 TMZ Resp/Non-Resp
Pair 2 TMZ Resp/Resp
Pair 3 TMZ Non-Resp/Resp

#### Figure S2

**A** GBM TMZ Non Responder (Pseudopalisade Structures)

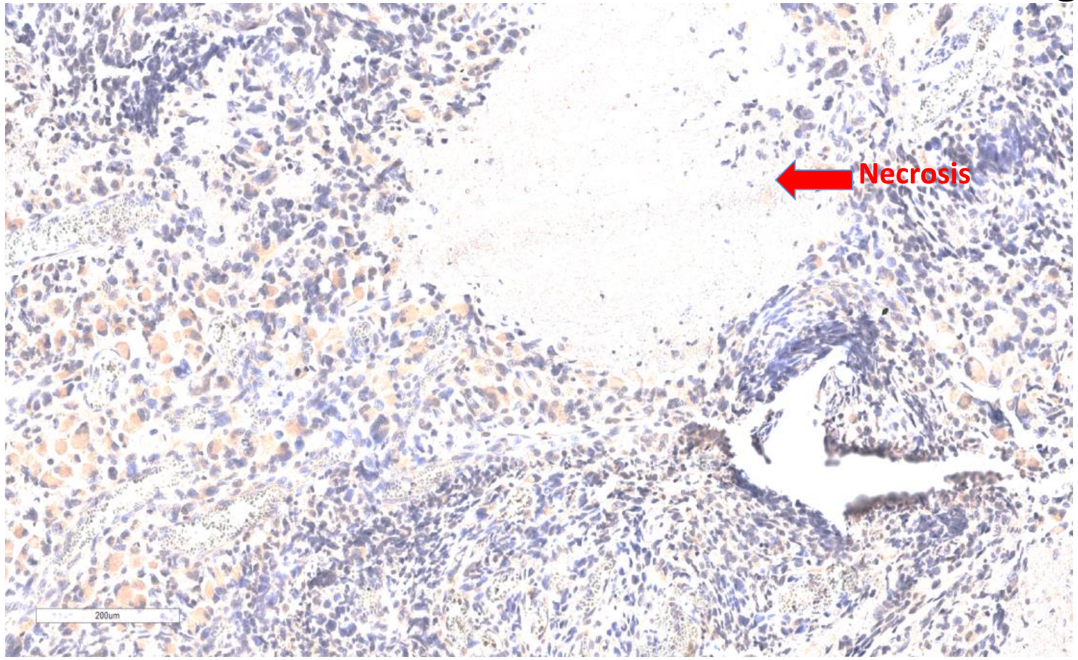


**B** GBM TMZ Responder (Pseudopalisade Structures)

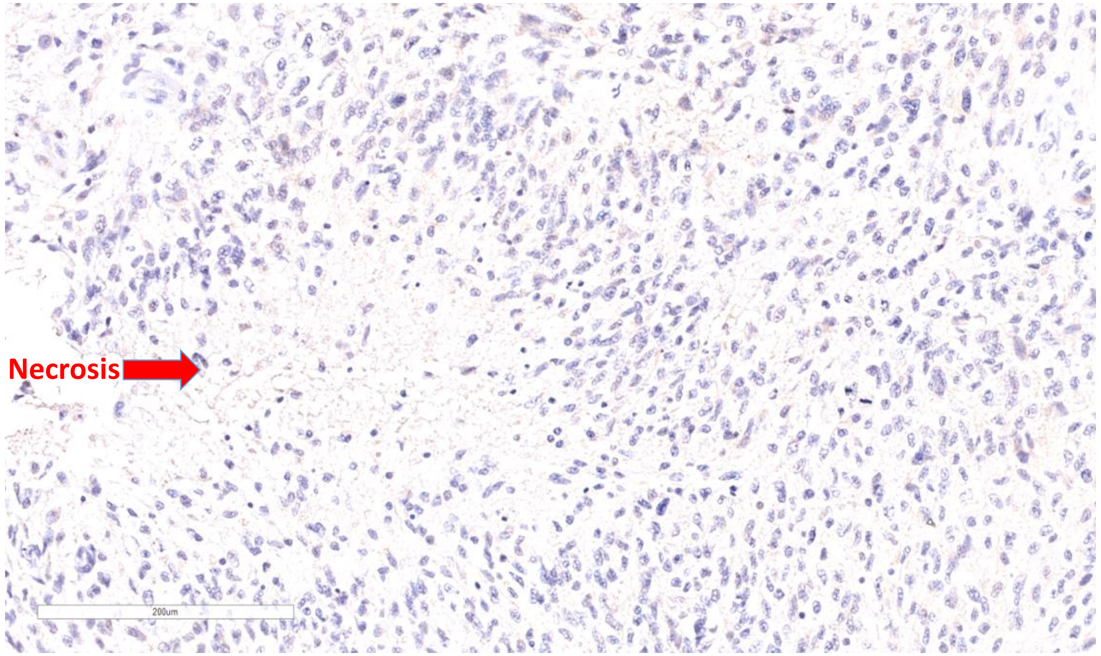


**C** GBM TMZ Non Responder (Tumor core)

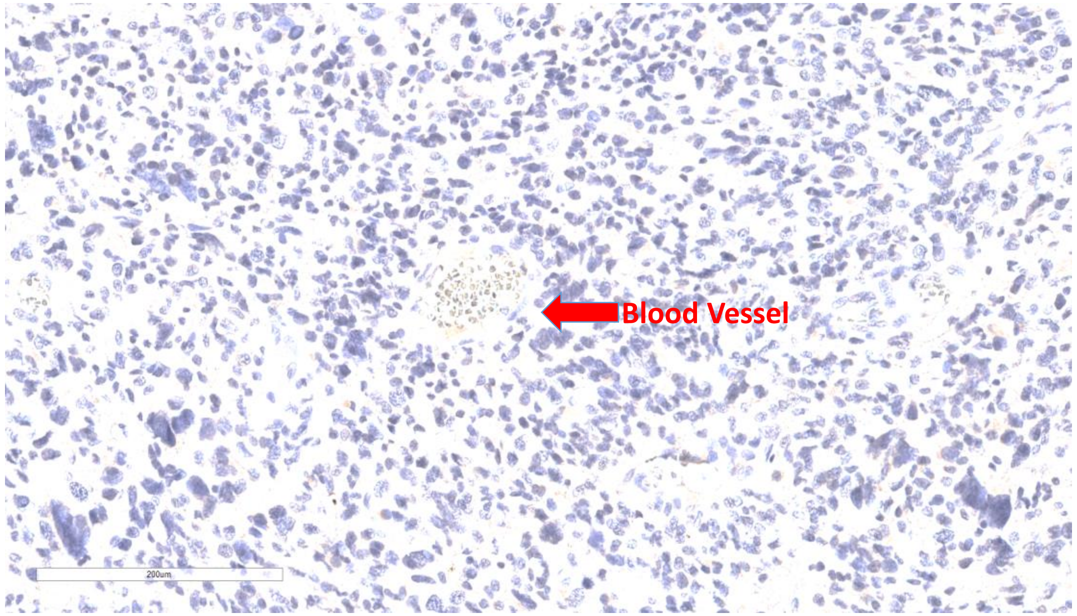


**D** GBM TMZ Responder (Tumor core)

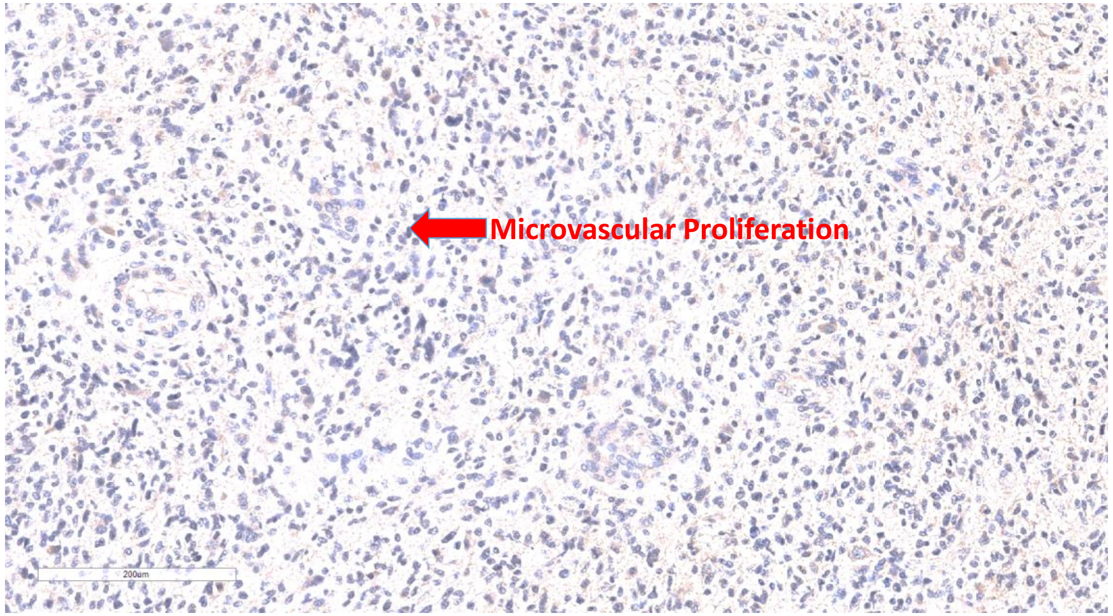

### Figure S3

A

OS survival of TCGA Agilent-4502A MGMT MEthyl

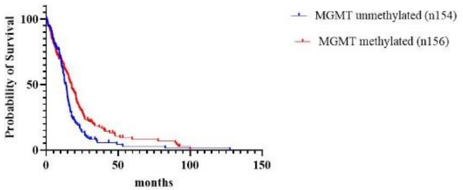


| | |
|---|---|
| P value | 0.0067 |
| P value | 0.0866 |

| | | |
|---|---|---|
| Median survival | | |
| MGMT unmethylated (n154) | 13.90 | |
| MGMT methylated (n156) | 17.60 | |
| Ratio (and its reciprocal) | 0.7898 | 1.266 |

B

OS survival of TCGA HG-UG133A MGMT methylation status

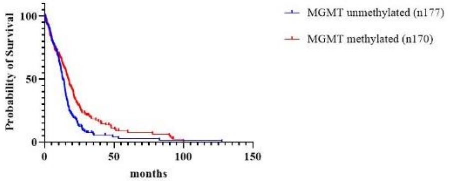


| | |
|---|---|
| P value | 0.0021 |
| P value | 0.0372 |

| | | |
|---|---|---|
| Median survival | | |
| MGMT unmethylated (n177) | 12.90 | |
| MGMT methylated (n170) | 17.00 | |
| Ratio (and its reciprocal) | 0.7588 | 1.318 |

C

OS survival of TCGA RNAseq MGMT methylation status

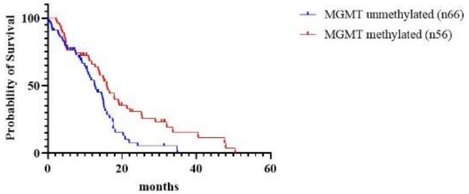


| | |
|---|---|
| P value | 0.0081 |
| P value | 0.0750 |

| | | |
|---|---|---|
| Median survival | | |
| MGMT unmethylated (n66) | 12.60 | |
| MGMT methylated (n56) | 15.90 | |
| Ratio (and its reciprocal) | 0.7925 | 1.262 |

D

OS survival of TCGA Agilent-4502A BIRC3 cutoff and MGMT methylation status

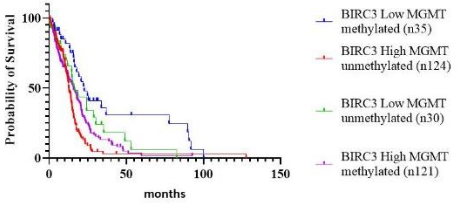


| | |
|---|---|
| P value | <0.0001 |

| | BIRC3 Law MGMT methylated (n35) | BIRC3 High MGMT unmethylated (n124) | BIRC3 Law MGMT unmethylated (n30) | BIRC3 High MGMT methylated (n121) |
|---|---|---|---|---|
| Median survival | 22.7 | 12.7 | 16.4 | 15.7 |

E

OS survival of TCGA HG-UG133A cutoff and MGMT methylation status

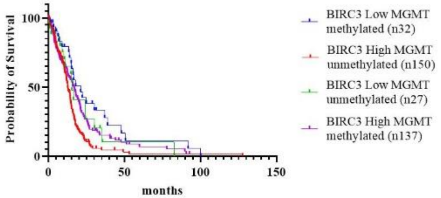


| | |
|---|---|
| P value | 0.0015 |

| | BIRC3 Low MGMT methylated (n32) | BIRC3 High MGMT unmethylated (n150) | BIRC3 Low MGMT unmethylated (n27) | BIRC3 High MGMT methylated (n137) |
|---|---|---|---|---|
| Median survival | 21.1 | 12.6 | 15.4 | 16.6 |

F

OS survival of TCGA RNAseq BIRC3 cutoff and MGMT methylation status

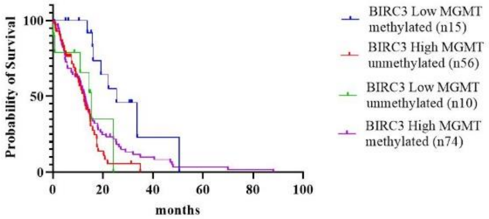


| | |
|---|---|
| P value | 0.0086 |

| | BIRC3 Low MGMT methylated (n15) | BIRC3 High MGMT unmethylated (n56) | BIRC3 Low MGMT unmethylated (n10) | BIRC3 High MGMT methylated (n74) |
|---|---|---|---|---|
| Median survival | 25.4 | 12.5 | 15.4 | 12.7 |

| Dependent: Surv(mytime, myoutcome) | | all | HR (univariable) | HR (multivariable) |
|---|---|---|---|---|
| MGMT_status | Methylated | 121 (50.0) | - | - |
| | Unmethylated | 121 (50.0) | 1.34 (1.04-1.73, p=0.026) | 1.34 (1.03-1.73, p=0.027) |
| cutoff | high | 195 (80.6) | - | - |
| | low | 47 (19.4) | 0.55 (0.39-0.76, p<0.001) | 0.55 (0.39-0.77, p<0.001) |

**Model Metrics:** Number in dataframe = 242, Number in model = 242, Missing = 0, Number of events = 242, Concordance = 0.560 (SE = 0.020), R-squared = 0.075( Max possible = 1.000), Likelihood ratio test = 18.866 (df = 2, p = 0.000)

##### Hazards Regression Plot

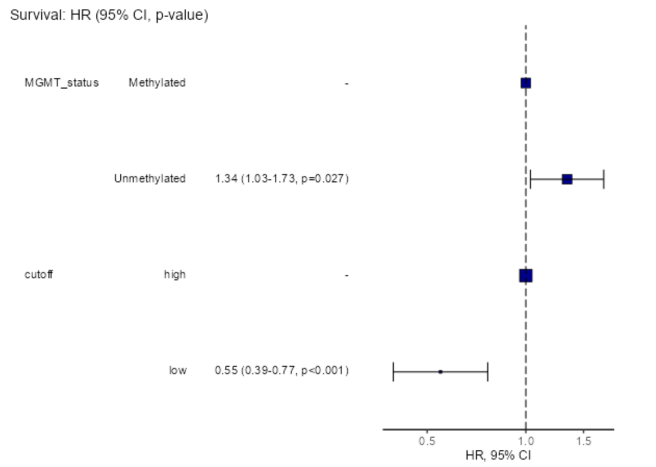


Agilent TCGA

| Dependent: Surv(mytime, myoutcome) | | all | HR (univariable) | HR (multivariable) |
|---|---|---|---|---|
| MGMT_status | Methylated | 56 (45.9) | - | - |
| | Unmethylated | 66 (54.1) | 1.80 (1.16-2.79, p=0.009) | 1.57 (1.01-2.45, p=0.047) |
| cutoff | high | 97 (79.5) | - | - |
| | low | 25 (20.5) | 0.41 (0.22-0.74, p=0.003) | 0.45 (0.25-0.83, p=0.010) |

**Model Metrics:** Number in dataframe = 122, Number in model = 122, Missing = 0, Number of events = 90, Concordance = 0.590 (SE = 0.032), R-squared = 0.113( Max possible = 0.996), Likelihood ratio test = 14.669 (df = 2, p = 0.001)

##### Hazards Regression Plot

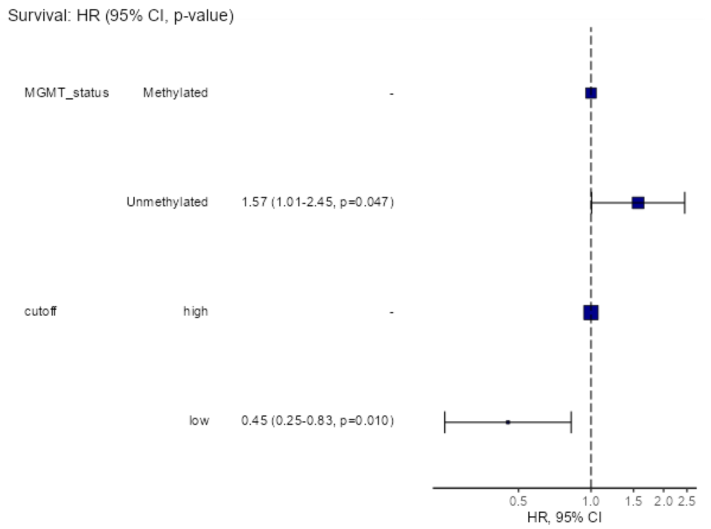


RNAseq

| Dependent: Surv(mytime, myoutcome) | | all | HR (univariable) | HR (multivariable) |
|---|---|---|---|---|
| MGMT_status | Methylated | 170 (49.0) | - | - |
| | Unmethylated | 177 (51.0) | 1.46 (1.15-1.86, p=0.002) | 1.44 (1.13-1.84, p=0.003) |
| mRNAcutoff | high | 288 (83.0) | - | - |
| | low | 59 (17.0) | 0.65 (0.46-0.91, p=0.013) | 0.67 (0.48-0.93, p=0.019) |

**Model Metrics:** Number in dataframe = 347, Number in model = 347, Missing = 0, Number of events = 273, Concordance = 0.556 (SE = 0.018), R-squared = 0.044( Max possible = 0.999), Likelihood ratio test = 15.476 (df = 2, p = 0.000)

##### Hazards Regression Plot

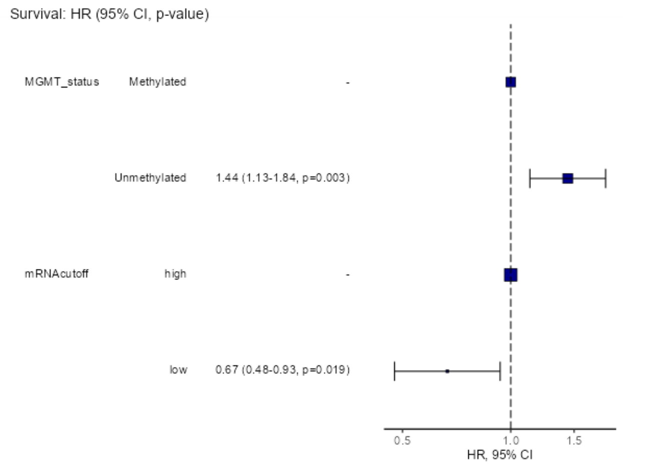


UGH133

**Figure S4**

A

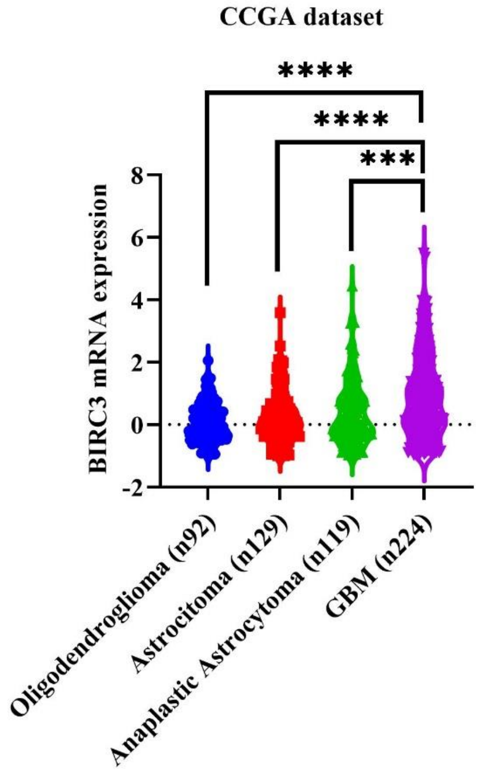


B

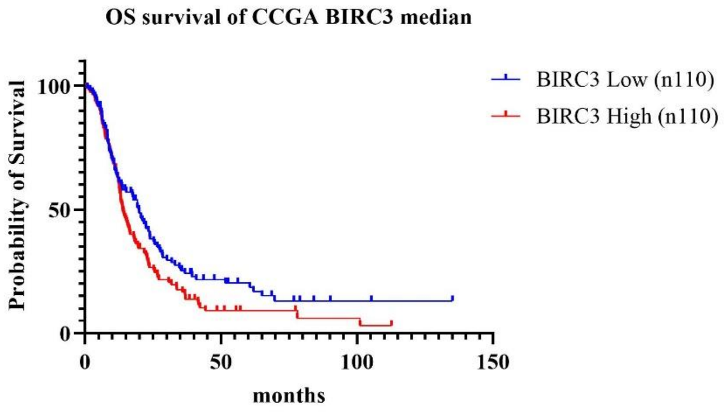


| | |
|---|---|
| P value | 0.0438 |

| | |
|---|---|
| P value | 0.1463 |

| | | |
|---|---|---|
| Median survival | | |
| BIRC3 Low (n110) | 19.80 | |
| BIRC3 High (n110) | 13.90 | |
| Ratio (and its reciprocal) | 1.424 | 0.7020 |
| 95% CI of ratio | 1.064 to 1.907 | 0.5244 to 0.9398 |

**Figure S5**

Figura S6

A

**OS and BIRC3 mRNA expression in Italian case study (primary tumors)**

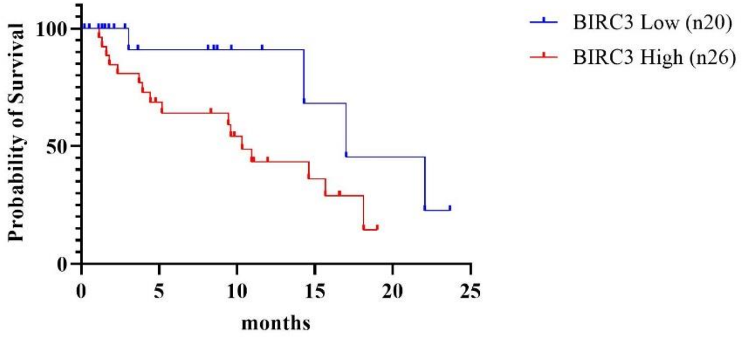


| | |
|---|---|
| P value | 0.0502 |
| P value | 0.0451 |

| | | |
|---|---|---|
| Median survival | | |
| BIRC3 Low (n20) | 17.00 | |
| BIRC3 High (n26) | 10.33 | |
| Ratio (and its reciprocal) | 1.645 | 0.6078 |
| 95% CI of ratio | 0.5500 to 4.921 | 0.2032 to 1.818 |

B

**PFS and BIRC3 mRNA expression in Italian case study (primary tumors)**

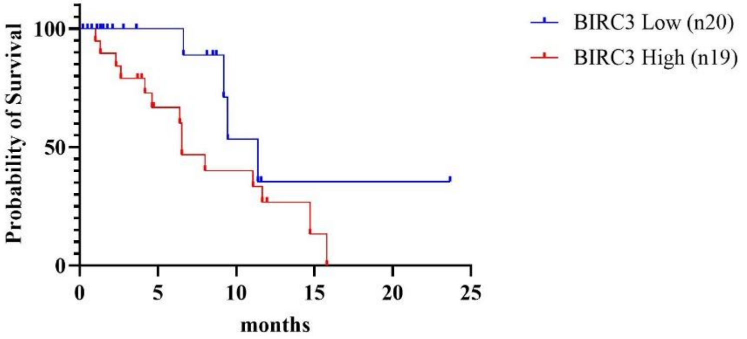


| | |
|---|---|
| P value | 0.0735 |
| P value | 0.0287 |

| | | |
|---|---|---|
| Median survival | | |
| BIRC3 Low (n20) | 11.40 | |
| BIRC3 High (n19) | 6.533 | |
| Ratio (and its reciprocal) | 1.745 | 0.5731 |
| 95% CI of ratio | 0.5743 to 5.301 | 0.1886 to 1.741 |

Figura S7

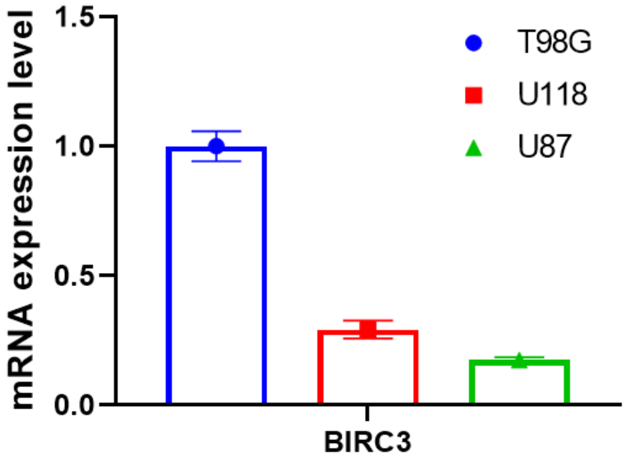
